# Experimental reproducibility limits the correlation between mRNA and protein abundances in tumour proteomic profiles

**DOI:** 10.1101/2021.09.22.461108

**Authors:** Swathi Ramachandra Upadhya, Colm J. Ryan

**Affiliations:** School of Computer Science and Systems Biology Ireland, University College Dublin, Dublin, Ireland

**Keywords:** proteomics, proteogenomics, machine-learning, reproducibility, transcriptomics, gene-expression, post-transcriptional regulation, cancer

## Abstract

Large-scale studies of human proteomes have revealed only a moderate correlation between mRNA and protein abundances. It is unclear to what extent this moderate correlation reflects post-transcriptional regulation and to what extent it reflects measurement error. Here, by analysing replicate profiles of tumours and cell lines, we show that there is considerable variation in the reproducibility of measurements of transcripts and proteins from individual genes. Proteins with more reproducible measurements tend to have higher mRNA-protein correlation, suggesting that measurement reproducibility accounts for a substantial fraction of the unexplained variation between mRNA and protein abundances. The reproducibility of individual proteins is somewhat consistent across studies and we exploit this to develop an aggregate reproducibility score that explains a substantial amount of the variation in mRNA-protein correlation across multiple studies. Finally, we show that pathways previously reported to have higher-than-average mRNA-protein correlation may simply contain members that can be more reproducibly quantified.

## Introduction

Proteins are the primary actors in our cells, responsible for almost all biological activities. Therefore understanding how protein abundances vary between healthy and disease states can provide an insight into how biological activities are altered in disease conditions. Among patients with the same disease, e.g. breast cancer, variation in protein abundances may explain differences in survival outcomes (Ősz et al., 2021) and drug responses (Shenoy et al., 2020). Consequently significant efforts have been made recently to characterise proteomes across large patient cohorts (Ellis et al., 2013). However, our ability to quantify protein abundances at scale has lagged behind our ability to sequence genomes and quantify mRNA abundances. Large-scale efforts to molecularly characterise healthy and disease samples from humans have therefore primarily focussed on DNA sequence variation and transcriptomic variation. For instance, the Cancer Genome Atlas project has quantified transcriptomes for ~1080 samples from breast tumours (Berger et al., 2018), while the proteomic equivalent, the NCI Clinical Proteomic Tumour Analysis Consortium (CPTAC) (Ellis et al., 2013), has used mass spectrometry (MS) proteomics to quantify ~105 breast tumour samples (Mertins et al., 2016). Alternative proteomic profiling approaches, such as reverse-phase protein arrays (RPPA), have been used to quantify much larger numbers of samples (Li et al., 2013) but these are limited to a small number (~200) of pre-specified proteins with available antibodies. For the unbiased quantification of thousands of proteins, mass spectrometry approaches remain the only viable option.

As transcriptomes are easier to quantify than proteomes, mRNA abundances are often used as a proxy for protein abundances. However, the relationship between mRNA abundances and protein abundances is complex, non-linear, and varies significantly from protein to protein. Consistent with this, large-scale studies in humans and model organisms have revealed that for most genes there is only a moderate correlation between mRNA and protein abundances (Buccitelli and Selbach, 2020; Vogel and Marcotte, 2012). We note that correlations between mRNA and protein abundances can be calculated in two different ways – across all proteins within a given sample (i.e. in a given cell line are the most abundant proteins also the most abundant transcripts?) or for a single protein across multiple samples (i.e. do the samples with the highest levels of a specific protein also have the highest number of transcripts coding for that protein?) (Franks et al., 2017; Liu et al., 2016; Vogel and Marcotte, 2012). Here, we are concerned with variation across individuals, and so throughout when we discuss mRNA-protein correlations we are calculating the correlation between the protein and transcript abundance for an *individual* protein *across* samples.

Tumour samples in particular have been subject to transcriptomic and proteomic profiling efforts and these have provided insight into how variation in mRNA abundances across individuals is associated with variation in protein abundances across the same individuals. These studies have reported an average mRNA-protein correlation in the range of ~0.2-0.5 (Mertins et al., 2016; Zhang et al., 2014, 2016). This moderate correlation between mRNA and protein abundances can be attributed to both biological and technical factors. Major biological factors that influence mRNA-protein correlation include translation rates that vary across proteins and conditions; highly variable half-lives for both proteins and mRNAs; and post-translational modifications that can alter protein stability and degradation (Buccitelli and Selbach, 2020).

Different proteins have been observed to have very different mRNA-protein correlations, and pathway enrichment analyses have identified specific functional groups with lower or higher than average mRNA-protein correlation. For instance, a number of metabolic pathways have been shown to have higher-than average mRNA-protein correlation (Clark et al., 2019; Huang et al., 2021; Jarnuczak et al., 2021; Mertins et al., 2016; Zhang et al., 2014, 2016), suggesting limited post-transcriptional regulation of these proteins. In contrast, subunits of large protein complexes have been shown to have lower than average mRNA-protein correlations, suggesting significant post transcriptional regulation. This can, at least partially, be attributed to the fact that proteins are often stabilised by their interactions with other complex members with subunits produced in excess being more rapidly degraded. A consequence of this is that proteins belonging to the same protein complex frequently display coordinated changes of abundance which are evident at the proteomic but not transcriptomic level (Gonçalves et al., 2017; Ryan et al., 2017; Taggart et al., 2020; Wang et al., 2017; Wu et al., 2013).

Our technical ability to accurately and reproducibly quantify both mRNAs and proteins is potentially a major factor that influences the mRNA-protein correlation. If the error in our measurements is large, we would expect this error to reduce the correlation between mRNA and protein even in the absence of the biological factors outlined above. A number of studies have separately assessed the reproducibility of either mRNA (’t Hoen et al., 2013; Marioni et al., 2008; SEQC/MAQC-III Consortium, 2014) or proteomic (Casey et al., 2017; Tabb et al., 2010) profiling approaches. Others have explored how measurement errors in mRNA or proteomic profiling can influence the reported correlation between mRNA and protein abundances. These have focussed on within sample correlations (across all proteins within a single sample / cell line) rather than across samples (for individual proteins across many samples) (Csárdi et al., 2015; Li et al., 2014).

Here, to understand the influence of measurement reproducibility on mRNA-protein correlation, we analyse studies of tumours and cancer cell lines with replicate proteomic profiles. We find that the correlation across replicate proteomes is not extremely high, with a median per-protein correlation of ~0.5 across replicates. This suggests that even in the absence of biological factors such as post-translational modifications and variable degradation we would not anticipate an mRNA-protein correlation anywhere close to 1. We find that some proteins are more reproducibly measured than others and that, in general, proteins that are reproducible in one study tend to be reproducible in others. We exploit this to create an ‘aggregate protein reproducibility’ ranking and show across multiple proteogenomic studies that proteins that are more reproducibly measured tend to have higher mRNA-protein correlations. We identify a number of factors, such as protein abundance, that influence protein measurement reproducibility. We also find that the reproducibility of mRNA measurements can, to some extent, independently explain the mRNA-protein correlations observed in proteogenomic studies. Finally we show that some pathways previously identified as having higher than average mRNA-protein correlation may simply result from the proteins in question being more reproducibly measured.

## Results

### A standardised pipeline reveals differences in the mRNA-protein correlation across studies

The average mRNA-protein correlation reported for different tumour proteomic profiling efforts varies substantially across studies – ranging from 0.23 in an early proteomic study of colorectal cancer (Zhang et al., 2014) to 0.53 in a recent study of lung adenocarcinoma (Gillette et al., 2020) (Table 1). However, it is not meaningful to directly compare the reported correlations because the methods used to quantify the mRNA-protein correlation have varied across studies – different studies have used different summary statistics (mean vs median), different correlation metrics (Pearson vs Spearman), and different criteria for protein inclusion (e.g. no missing values, at least 30% measured values, only the 10% most variable proteins) (Table 1). To enable a more direct comparison across studies, we calculated the mRNA-protein correlation for thirteen proteomic studies using a standardised pipeline. The datasets analysed comprise ten studies of tumour samples (Clark et al., 2019; Dou et al., 2020a; Gillette et al., 2020; Huang et al., 2021; Krug et al., 2020; Mertins et al., 2016; Vasaikar et al., 2019; Wang et al., 2021; Zhang et al., 2014, 2016), two studies of cancer cell lines (Guo et al., 2019; Nusinow et al., 2020), and one study of healthy tissues (Jiang et al., 2020). Within each study, we calculated the median Spearman’s correlation between mRNA and protein for all proteins that were measured in at least 80% of samples (Methods; Table 1 and S1). Across all studies, the median recalculated correlation was 0.43 (BrCa 2020 (Krug et al., 2020)) with a maximum of 0.55 (LUAD (Gillette et al., 2020)) and a minimum of 0.21 (CRC (Zhang et al., 2014)). In some instances the recalculated correlation was similar to that originally reported but in others there was a substantial difference. For example, the correlation recalculated for endometrial cancer (0.48) was the same as originally reported (Dou et al., 2020b) while the recalculated correlation for colon cancer was much lower than that reported by the authors (0.27 vs 0.48) (Vasaikar et al., 2019). This large difference is because the colon cancer study originally reported the mean mRNA-protein correlation for only the 10% most variable proteins, rather than the full set of proteins. These highly variable proteins have higher than average mRNA-protein correlation.

**Table 1.** Analysis of mRNA-protein correlation using a standardised pipeline.

| Data | Published Year | Reported Correlation | Protein inclusion criterion in reported correlation | Computed median Spearman Correlation |
| --- | --- | --- | --- | --- |
| GTEX 32 Healthy Tissues (GTEX) | 2020 | 0.46 | < 5 tissues with missing values for both protein and RNA measurements | 0.51 |
| Cancer cell line encyclopaedia (CCLE) | 2020 | 0.48 | Quantified in at least one ten-plex (9 cell lines) | 0.46 |
| NCI-60 cancer cell lines (NCI60) | 2019 | Not reported | - | 0.36 |
| Glioblastoma (GBM) | 2021 | Not reported | - | 0.50 |
| Head and Neck Squamous Cell carcinoma (HNSCC) | 2021 | 0.52 | < 50% missing values | 0.54 |
| Lung adenocarcinoma (LUAD) | 2020 | 0.53 | < 50% missing values | 0.55 |
| Endometrial Cancer (EC) | 2020 | 0.48 | Contain mRNA and protein measurements across all patients | 0.48 |
| Breast cancer (BrCa 2020) | 2020 | 0.41 | Contain mRNA and protein measurements (proteins < 70% missing values) | 0.44 |
| Clear cell renal carcinoma (ccRCC) | 2019 | 0.43 | Contain mRNA and protein measurements across all patients | 0.41 |
| Colon Cancer (Colon) | 2019 | 0.48 | top 10% most variably expressed genes quantified in both platforms | 0.27 |
| Ovarian cancer (Ovarian) | 2016 | 0.45 | Contain mRNA and protein measurements across all patients | 0.41 |
| Breast cancer (BrCa 2016) | 2016 | 0.39 | Contain mRNA and protein measurements across all patients passing quality control checks. | 0.42 |
| Colon and rectal cancer (CRC 2014) | 2014 | 0.23 | Protein measurement with average spectral count across all patients $\geq 1.4$ | 0.21 |

More recent studies appear to have higher mRNA-protein correlations – e.g. we observe a mean of 0.49 for studies published after 2019 vs 0.35 for studies published in 2016 or earlier (Table 1). This cannot simply be attributed to differences in the cancer types studied in different years, as the two cancer types profiled twice (colon and breast) see an improvement from the earlier studies (Table 1). This would suggest that technical and experimental factors may influence the reported mRNA-protein correlations and that improvements in either technology or experimental protocols have resulted in improved mRNA-protein correlations over time.

### The correlation across replicate proteomic profiles is only moderate

To assess the reproducibility of mass spectrometry based proteomic measurements, we analysed three studies containing replicate proteomic profiles – ovarian tumour samples (Zhang et al., 2016), colon tumour samples (Vasaikar et al., 2019) and cancer cell lines of mixed lineages from the Cancer Cell Line Encyclopedia (CCLE) (Nusinow et al., 2020) (Figure 1A). The nature of the replicates varies across the different studies – for ovarian cancer the same tumour sample was profiled in two different laboratories, for the cancer cell lines biological replicates were performed within the same lab one year apart, while for colon cancer the same tumour samples were profiled with two different MS techniques, i.e., isotope based protein quantification (TMT-10) and label-free spectral counting MS. Thus there is diversity in the replicate proteomic profiles in terms of sample types (tumour samples and cancer cell lines), sites, and techniques used to quantify the proteins.

**Figure 1.**
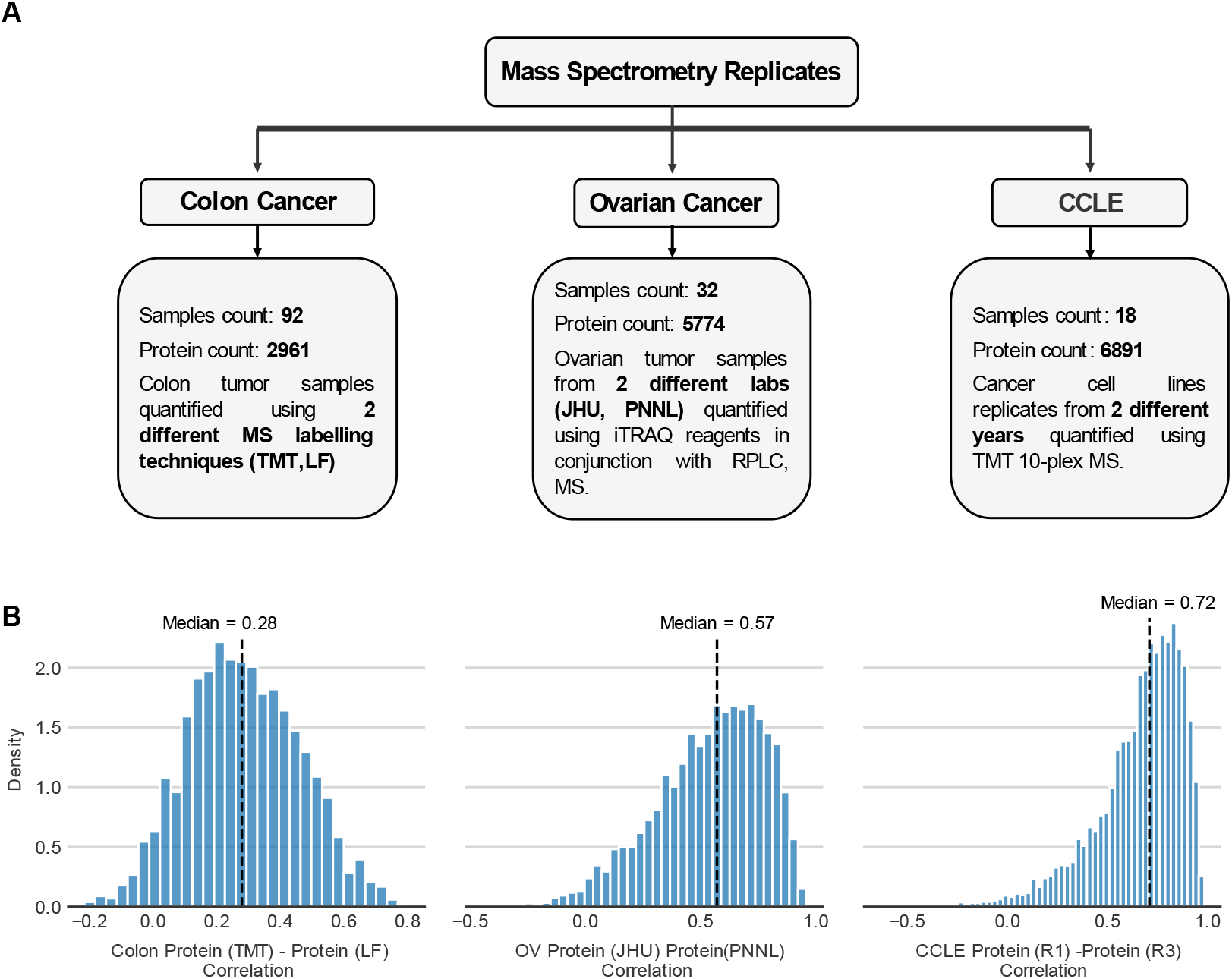
Protein-protein reproducibility across replicates are moderate and variable. (A) Overview of the replicates available for the three different proteomic studies (B) For each study, we calculate the correlation for individual proteins across the proteomic replicates. The distribution of the protein-protein reproducibility is shown in the histogram for all measured proteins. For each study, the black dotted line represents the median.

Many biological factors that influence mRNA-protein correlation, such as post-transcriptional regulation, are not relevant in the case of replicate measurements of proteins and so we expected the replicate proteomic profiles to be more highly correlated than mRNA and protein profiles. This was indeed the case for all studies. The median protein-protein reproducibility for the replicate proteomic profiles from the CCLE dataset was 0.72 (Figure 1B; Table S2) whereas the median mRNA-protein correlation was only 0.48 (Table 1). The median proteinprotein reproducibility for the replicate proteomic profiles of ovarian tumours was 0.57 (Figure 1B) which is higher than the median mRNA-protein correlation of 0.41 (Table 1). The replicate protein-protein reproducibility for the colon study (median 0.28) was much lower than that observed for the other studies. However, it was still higher than the median calculated mRNA-protein correlation (0.21). One reason for the colon study to have a low median protein-protein reproducibility is that one of the two replicate proteomic profiles is quantified using label-free/spectral counting mass spectrometry which is not as accurate as the stable isotop-based protein quantification methods (Liu et al., 2016). Overall, we can conclude that although protein-protein reproducibility is consistently higher than mRNA-protein correlations, the protein-protein reproducibility is still only moderate.

### Proteins with higher reproducibility have higher mRNA-protein correlation

The moderate correlations reported between mRNA and protein abundances have been attributed to a variety of biological factors, including post transcriptional regulation, varying translation rates and varying degradation rates (Buccitelli and Selbach, 2020; Payne, 2015; Vogel and Marcotte, 2012). However, our observation that some proteins can be quantified more reproducibly than others suggests that noise in quantification may also be a major factor. If this is the case we would expect that proteins that can be more reproducibly quantified will have a higher mRNA-protein correlation. To assess this, for each study we used the replicate proteomic profiles to stratify the proteins into deciles, ranging from the 10% of proteins with the lowest protein-protein reproducibility to the 10% with the highest protein-protein reproducibility (Methods). We then calculated the mRNA-protein correlation for all of the proteins within each decile. We found, for all three studies, that the median mRNA-protein correlation increases with protein-protein reproducibility (Figure 2). The colon cancer study shows a difference in the median mRNA-protein correlation of 0.33 between the first and last deciles of protein reproducibility. Similarly, ovarian cancer data shows a difference of 0.35 and the CCLE data shows a difference of 0.37. This indicates that the reproducibility of proteomic measurements has a major impact on the calculated mRNA-protein correlation. We used a linear regression model to understand how much of the variation in mRNA-protein correlation can be explained by variation in protein-protein reproducibility and found that it explains approximately 14%, 17% and 23% in the ovarian, CCLE, and colon studies respectively (Methods; Figure 2 and S1A).

**Figure 2.**
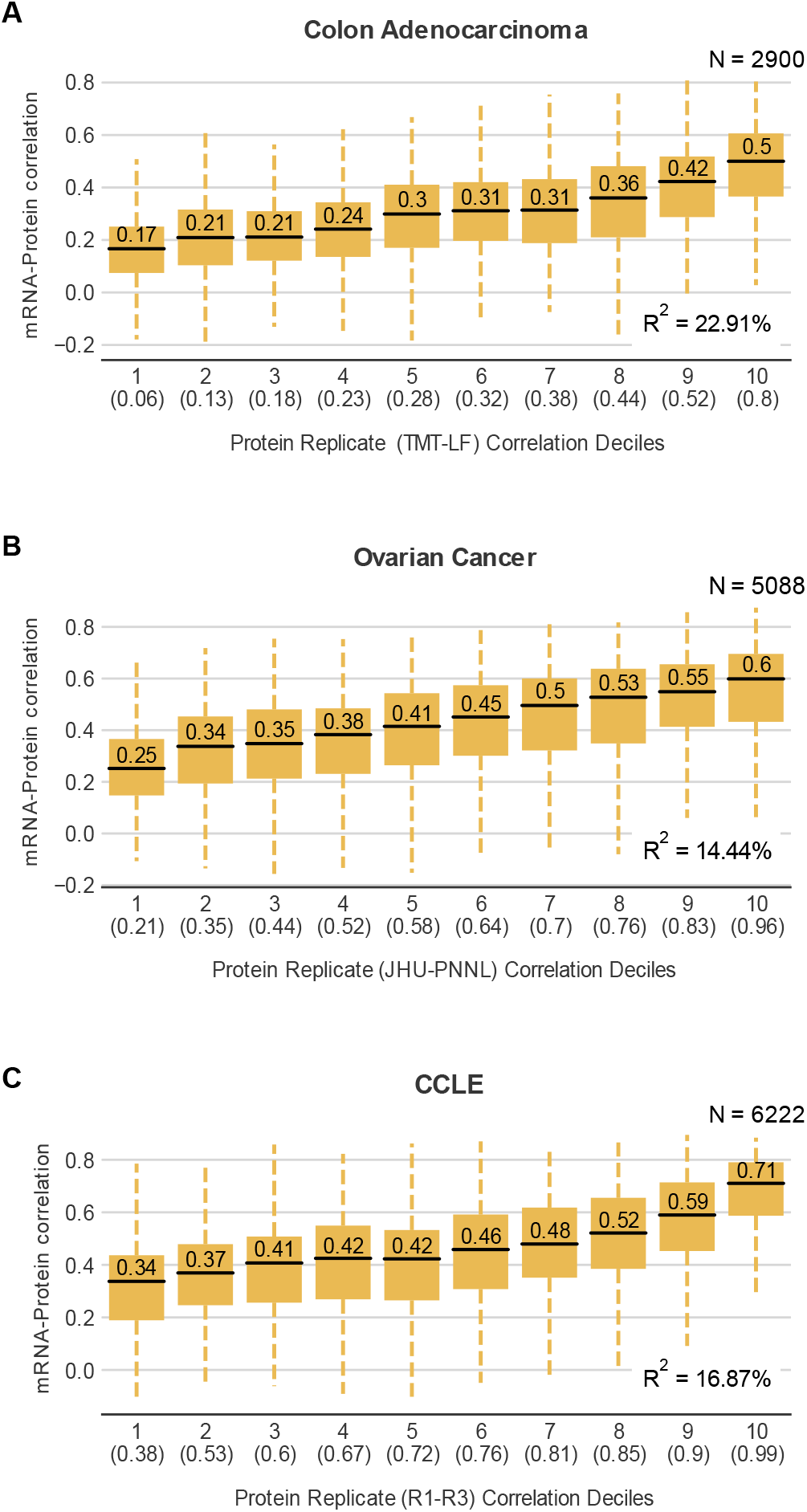
Proteins with higher reproducibility have higher mRNA-protein correlation. Boxplots showing the distribution of mRNA-protein correlation for proteins binned according to their protein-protein reproducibility in the colon (A), ovarian (B) and CCLE studies (C). The total number of proteins considered for each plot is indicated at the top right corner. The bins are deciles – each containing ~10% of the proteins. The decile is indicated on the X-axis along with the highest correlation between experimental replicates present within that decile. For each box plot, the black central line represents the median, the top and bottom lines represent the 1st and 3rd quartile, and the whiskers extend to 1.5 times the interquartile range past the box. Outliers are not shown. The median of each decile is indicated above the black central line for each box plot. The R^2^ obtained from regressing the mRNA-protein correlation on protein-protein reproducibility is in the bottom-right corner.

Previous work has identified protein complex membership as the factor most predictive of variation in mRNA-protein correlation, with subunits of protein complexes typically having lower than average mRNA-protein correlation (Gonçalves et al., 2017; Ryan et al., 2017). Using the same linear modelling approach as above we found that protein complex membership explains approximately 3%, 8%, 6.7% of the variation in the ovarian, CCLE, and colon studies respectively (Figure S1A). This suggests that noise in the quantification of protein abundances explains much more (on average ~3 times) of the variance in the mRNA-protein correlation than the most predictive previously identified factor. Combined, the protein-protein reproducibility and protein complex membership features explained approximately 17%, 23%, 26% of the variation in mRNA-protein correlation in the ovarian, CCLE and colon studies respectively (Figure S1A). This is significantly more than protein complex membership or protein-protein reproducibility alone (p-value < 0.001, Likelihood ratio test) suggesting that protein complex membership and protein reproducibility independently contribute to the variation in mRNA-protein correlation. This is evident when binning proteins into reproducibility deciles – although proteins which are complex subunits are present in every decile, they have consistently lower mRNA-protein correlations (Figure S1B-D).

### Proteins with high reproducibility in one study are also highly reproducible in other studies

In addition to providing a summary of how reproducible the protein measurements from each study are on average, the replicate profiles enable us to see which proteins are most reproducibly quantified overall. In the CCLE study the median correlation between replicate measurements calculated across all proteins was 0.72 but this ranged from −0.2 to 1.0 for individual proteins. Similarly the median for all proteins in the ovarian study was 0.57 but the individual correlations ranged from −0.6 to 1.0, and the median for the colon tumour study was 0.28 with a range from −0.2 to 0.8. This suggests that, at least within individual studies, some proteins may be more reproducibly quantified than others.

To understand whether the same proteins were reproducibly quantified across multiple studies, we analysed pairs of studies together. We found that there was a moderate correlation (0.38) between the protein reproducibility calculated using the ovarian tumour replicates and the colon cancer replicates (Figure 3A). Combinations of other pairs of studies revealed similar moderate correlations – colon and CCLE (0.31); ovarian and CCLE (0.24) (Figure 3B-C). Although the nature of the samples (tumour versus cell line) and the quantification approaches (TMT / Label Free Quantification) varied across studies, this suggests that there is some agreement in terms of which proteins can be reproducibly quantified. In general, proteins that are highly reproducible in one study tend to be highly reproducible in others while proteins that show poor reproducibility in one study tend to show poor reproducibility in others (Figure 3).

**Figure 3.**
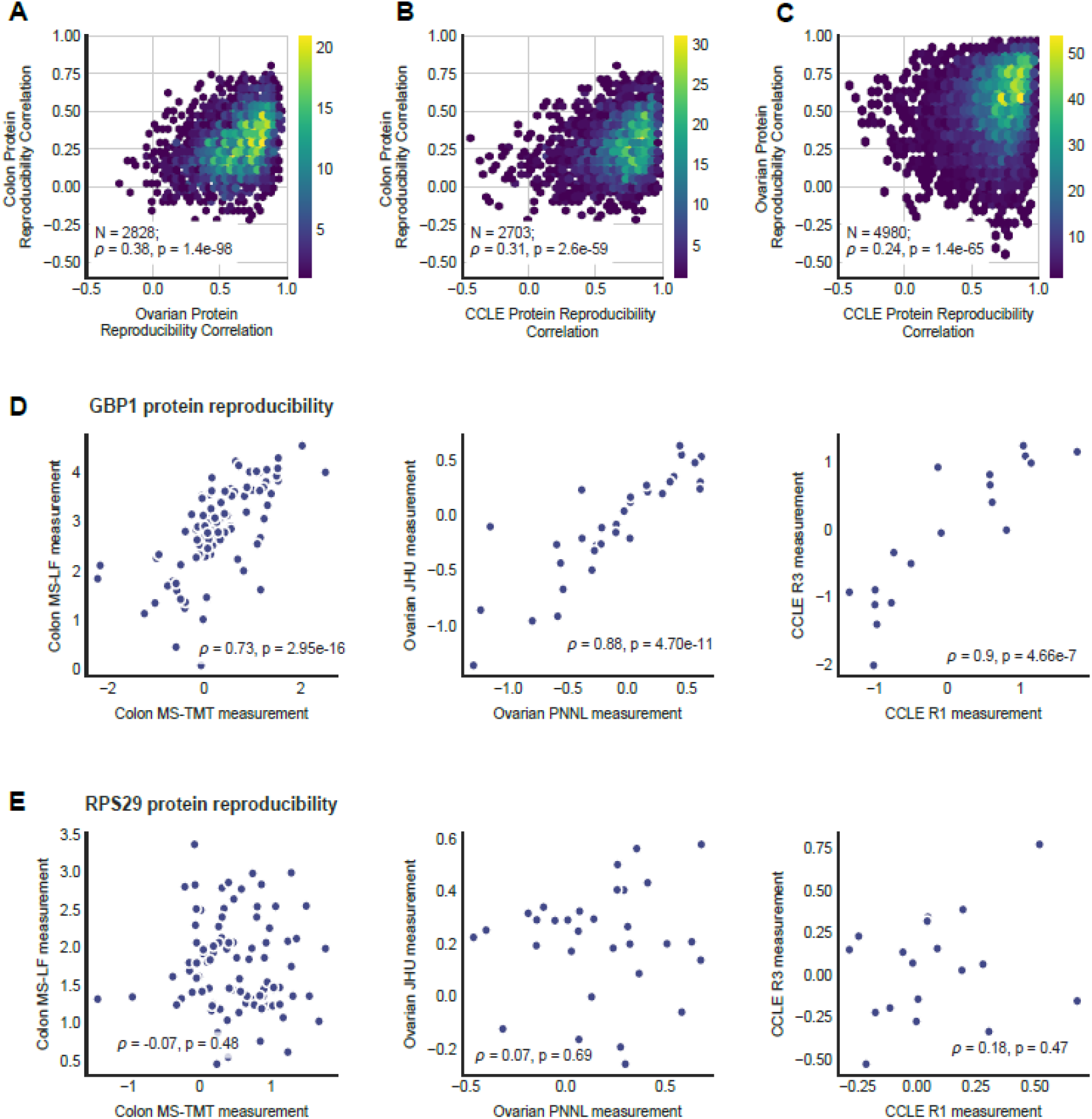
Proteins with high reproducibility in one study are also highly reproducible in other studies. (A-C) Binned heat maps showing the relationship between the protein-protein reproducibility calculated in different studies. Each heatmap shows the relationship between two studies, indicated on the x and y axes. The regions of the heatmaps are coloured according to the number of proteins present in the region as indicated in the colour bar. The number of proteins in common, and Spearman correlation between the two studies, with the associated p-value, are specified in the box for each of the plots. (D-E) For each study with experimental protein replicates, scatter plots illustrating the relationship between protein-protein reproducibility is shown for a protein with high reproducibility - GBP1 (D) and a protein with low reproducibility RPS29 (E). For each scatter plot, the Spearman correlation coefficient of the protein-protein reproducibility and the associated p-value is indicated at the bottom.

For example, GBP1 is one of the proteins with reproducibility that is consistently high across all three studies (Figure 3D) while RPS29 has consistent low reproducibility (Figure 3E).

### An integrated ranking of protein reproducibility partially explains the variable mRNA-protein correlation in 10 additional studies

Proteogenomic studies with large numbers of replicates, such as the three we analysed above, are the exception rather than the rule. Consequently for most studies we do not know how reproducible the proteomic measurements are. However, as noted above, proteins that are highly reproducibly quantified in one study are more likely to be highly reproducible in others. We therefore sought to aggregate the replicate protein correlations from all three studies (CCLE, ovarian, colon) into a single list containing a ranking of protein reproducibility (Methods; Figure S2A; Table S2). We evaluated a number of different aggregation approaches and found that a simple method using average normalized rank explained the most variance in the mRNA-protein correlation of the three studies containing proteomic replicates (Methods; Figure S2B). We used this approach to create a ranked order of protein reproducibility for the 5,211 proteins that were quantified in at least two out of the three studies. We then used this aggregated list to assess the extent to which ‘average’ protein reproducibility explains the varying mRNA-protein correlations observed in ten other studies: colorectal cancer (Zhang et al., 2014), clear cell renal carcinoma (Clark et al., 2019), endometrial cancer (Dou et al., 2020b), lung adenocarcinoma (Gillette et al., 2020), head and neck squamous cell carcinoma (Huang et al., 2021), glioblastoma (Wang et al., 2021), two breast cancer studies (Krug et al., 2020; Mertins et al., 2016), NCI-60 cancer cell lines (Guo et al., 2019) and healthy tissues from the GTEx project (Jiang et al., 2020) (Figure 4). For all these studies, we find that, in accordance with our previous observations – proteins with more reproducible measurements tend to have higher mRNA-protein correlations. Although the aggregated ranks are based on data from cancer studies we observe the same trend in healthy tissues obtained from the GTEx project (Figure 4I). Similarly, although the aggregated ranks are generated using studies that quantify proteins through data dependent acquisition (DDA) approaches, we observed the same trend for a study that quantified proteins using data independent acquisition (DIA) based proteomics (SWATH-MS) in the NCI-60 cancer cell lines (Figure 4J). In general, the mRNA-protein correlation increases with protein reproducibility for samples from both healthy and diseased conditions, and irrespective of the proteomic quantification approach.

**Figure 4.**
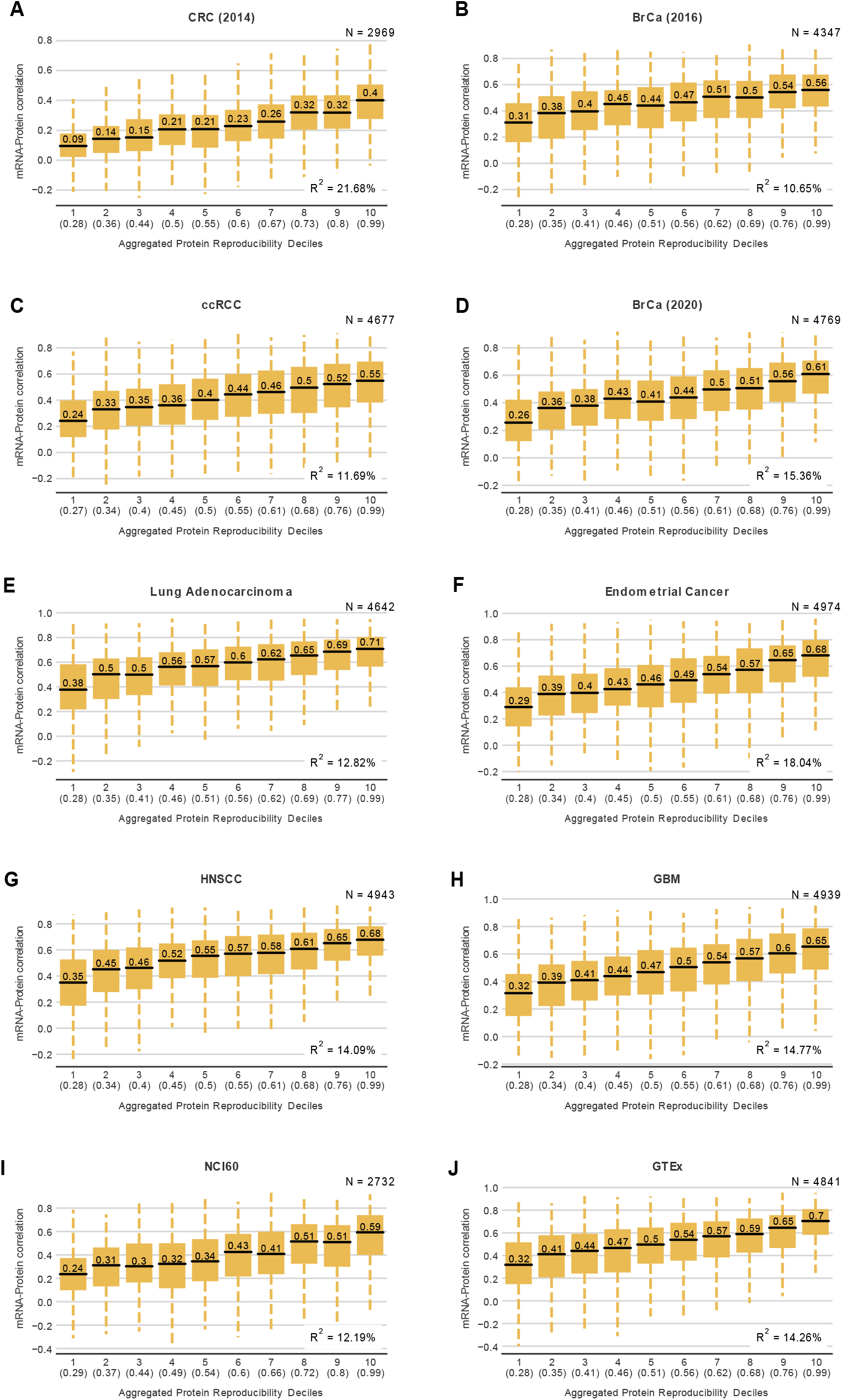
Aggregated protein reproducibility ranks partially explains the variable mRNA-protein correlation in 10 additional studies. (A-J) For studies without experimental proteomic replicates, boxplots showing the distributions of mRNA-protein correlation for proteins in each decile of the aggregated protein reproducibility ranks. (A-H) are the CPTAC tumour studies; (I) is the healthy tissues study from the GTEx Consortium and (J) is the NCI-60 cancer cell lines study wherein protein quantification, used for computing the mRNA-protein correlation, is obtained from data-independent acquisition based untargeted proteomics (SWATH-MS). The total number of proteins considered for each plot is in the top-right corner. The decile is indicated on the X-axis along with the highest score of the aggregated protein reproducibility rank present within that decile. For each box plot, the black central line represents the median, the top and bottom lines represent the 1st and 3rd quartile, and the whiskers extend to 1.5 times the interquartile range past the box. Outliers are not shown. The median of each decile is indicated above the black central line for each box plot. The R^2^ obtained from regressing the mRNA-protein correlation using the aggregated reproducibility ranks as a factor is in the bottom-right corner.

To quantify the amount of variation in mRNA-protein correlation that could be explained by our aggregated protein reproducibility ranks we used a linear regression model for the ten different studies. We found that the aggregated ranks explain ~10-20% (median 14%) of the variation in these studies (Figure 4).

To test if there was an advantage to using the aggregate protein reproducibility over protein reproducibility measured in either of the three individual studies (CCLE, ovarian, colon), we compared the variance explained by the aggregate ranks to that explained by each individual study. In all the ten studies without proteomic replicates, the aggregated ranks explained the variation in mRNA-protein correlation better than the ranks from any individual dataset (Figure S3).

Since there are orders of magnitude more tumour samples with quantified transcriptomes than proteomes, a number of efforts have been made to use machine learning to predict protein abundances from mRNA abundances (Fortelny et al., 2017; Li et al., 2019; Yang et al., 2020). Recently the NCI-CPTAC DREAM proteogenomics challenge engaged the community to predict protein abundances of tumour profiles using their corresponding genomic and transcriptomic information (Yang et al., 2020). The genomic, transcriptomic and proteomic data from CPTAC breast (Mertins et al., 2016) and ovarian (Zhang et al., 2016) tumour samples and their confirmatory studies were used for training and testing the machine learning models respectively. The best performing model for predicting protein abundances from mRNA abundances achieved an average correlation of 0.51 between actual and predicted values for breast cancer and 0.53 for ovarian cancer. Protein complex membership, protein half-lives and mRNA abundance were some of the factors identified that influence the predictability of proteins (Yang et al., 2020). We hypothesized that proteins whose measurements are highly reproducible could be predicted better using machine learning algorithms. Hence, we analysed the prediction scores from the best performing model using the protein reproducibility data. We observed a stark difference in the prediction scores of the lowest and highest deciles of the protein reproducibility (Figure S4A-B). While the lowest decile has a correlation of ~0.3 between the measurements and predictions, the highest decile has a correlation of ~0.7. On performing linear regression, we noticed that the protein-protein reproducibility measured using the ovarian, CCLE and Colon studies could explain approximately 12%, 13% and 15% of the variation in the prediction scores for breast cancer study, and 17%, 12% and 14% in ovarian cancer study respectively (Figure S4C). Notably, the aggregated ranks again outperformed the individual replicate correlations and could explain ~25% and 26% of the variation in the prediction scores of breast and ovarian cancer studies respectively (Figure S4C).

### Protein measurement reproducibility is influenced by abundance, variance and unique peptides

To understand what causes some proteins to be more reproducibly measured than others we analysed a number of factors that we hypothesised might influence the reliability of their measurements.

All of the studies analysed here make use of ‘bottom-up’ q uantification approaches where proteins are first digested into peptides; these peptides are then quantified using a mass spectrometer, and peptide quantifications converted into protein abundances computationally. This quantification is a stochastic process and there is no guarantee that every peptide in a given sample will be detected by the mass spectrometer. The quantification of proteins that have low abundance, and hence fewer detectable peptides, is especially likely to be subject to substantial stochastic variation. A small number of peptides missed can make a big difference to the quantification of these low abundance proteins, while for highly abundant proteins a few extra or missing peptides will make only a small difference. To assess the contribution of protein abundance to protein measurement reproducibility, we obtained the protein abundances measured in 201 tissue samples from 32 healthy human tissues collected by the GTEx project (Jiang et al., 2020). For each protein we calculated the mean abundance across all samples and tissues. We found a clear relationship between the mean protein abundance and the aggregated protein reproducibility rank – more abundant proteins are more reproducibly measured (Figure 5A). We performed similar analysis for the three individual proteomic replicate studies and found the result to be consistent (Figure S5A-C).

**Figure 5.**
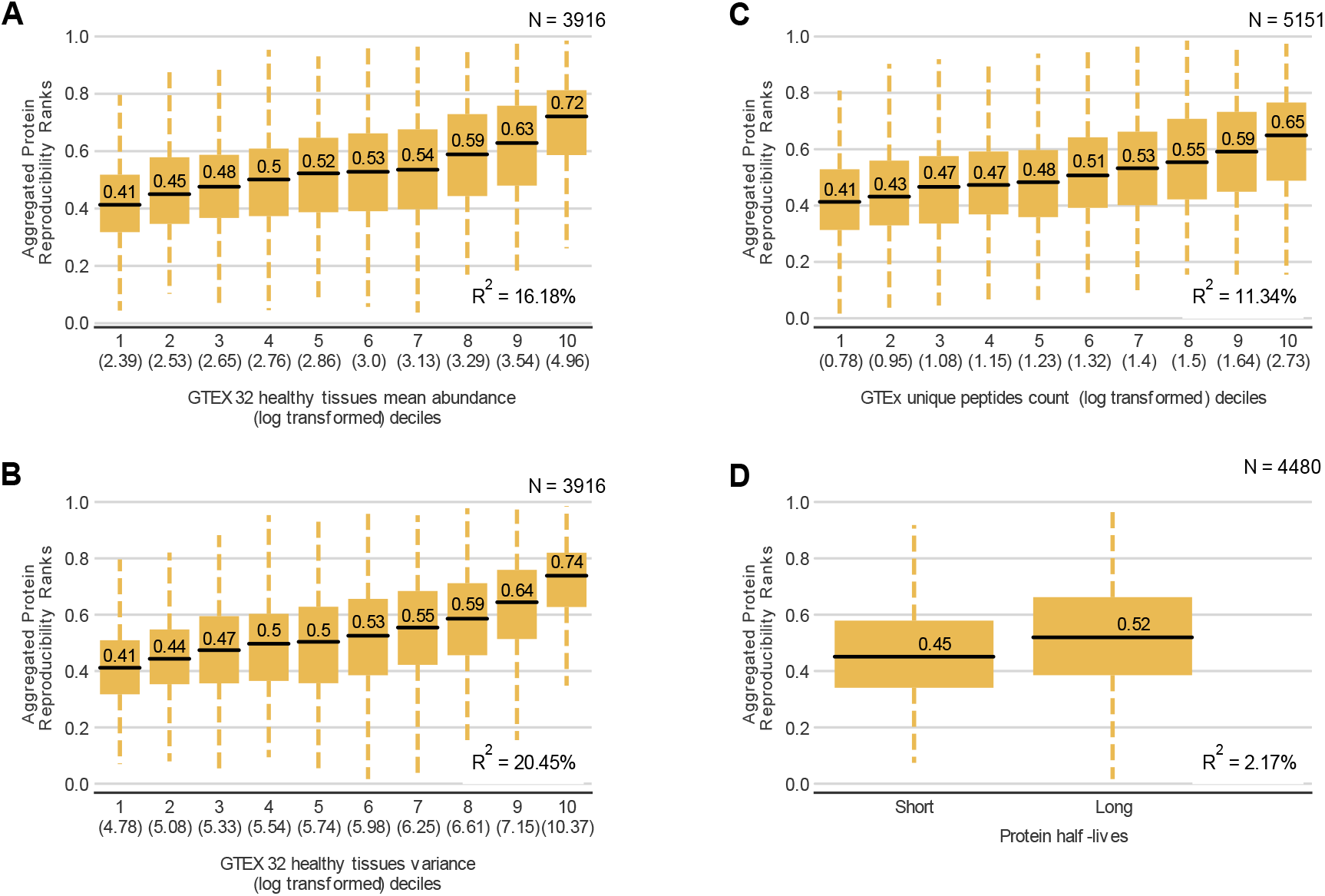
Protein reproducibility is mainly influenced by abundance, variance, unique peptides and not protein half-lives. Boxplots showing the distribution of aggregated protein reproducibility ranks for proteins binned according to protein abundance (A), variance (B) and number of unique peptides (C). The bins are deciles – each containing ~10% of the proteins. The decile is indicated on the X-axis along with the highest correlation between experimental replicates present within that decile. For each box plot, the black central line represents the median, the top and bottom lines represent the 1st and 3rd quartile, and the whiskers extend to 1.5 times the interquartile range past the box. Outliers are not shown. The median of each decile is indicated above the black central line for each box plot. (D) Boxplot showing the distribution of aggregated protein reproducibility ranks for proteins with short and long protein half-lives. For each plot, the R^2^ obtained from regressing the aggregated protein reproducibility ranks on each of the potential factors of protein reproducibility is in the bottom-right corner.

Protein whose abundances do not vary significantly across individuals are unlikely to have high mRNA-protein correlations, as correlation measures are dependent on their being meaningful variation in the data. Furthermore, as the variation observed experimentally is a likely a combination of both real biological variation and experimental noise, proteins with lower biological variation in abundance tend to be more affected by measurement noise. For each protein we computed the variance in protein abundance across samples from the GTEx project (Jiang et al., 2020). We then assessed the influence of this variance on the reproducibility of measurements of individual proteins. Similar to the mean protein abundance above, we found that proteins with a higher variance of protein abundance are more reproducibly measured (Figure 5B). Furthermore, the variance of protein abundance explains ~20% of the variation in the aggregated protein reproducibility ranks. This result was also true for the three individual proteomic replicate studies (Figure S5A-C).

The number of unique peptides generated per protein is also crucial for protein quantification by mass spectrometry. To assess the impact of this we identified the number of unique peptides identified per protein using the GTEx study. We stratified all proteins into deciles based on the number of unique peptides identified and found that the aggregated protein reproducibility increased with every decile of unique peptides identified (Figure 5C). This pattern was also evident in the protein reproducibility measured in each of the three individual studies (Figure S5A-C). Thus, the more unique peptides identified per protein, the higher the confidence of the measured protein levels.

One of the biological reasons proposed for the weak mRNA-protein correlation is the difference in mRNA and protein half-lives (Vogel and Marcotte, 2012). mRNAs typically have a half-life of 2.6-7 hours while proteins have half-lives ranging from a few seconds to a few days (Vogel and Marcotte, 2012). Recently, proteins with longer half-lives were found to be more predictable using machine learning irrespective of the transcript half-lives (Yang et al., 2020). As noted earlier, proteins with higher reproducibility are better predicted. This led us to assess protein half-life as a potential factor for the reproducibility of protein measurements. We obtained protein half-lives estimations from a previous publication (Zecha et al., 2018) and divided them into two categories - long and short half-lives (Methods) - as was done in (Yang et al., 2020). Although both categories of protein half-lives contain proteins with reproducibility scores ranging between 0 and 1, proteins with a long half-life have a better median protein reproducibility score (p-value = 9.70e-25, Mann-Whitney U test, two-sided; Figure 5D; Figure S5A-C).

We note that there is some correlation between the attributes considered, in particular more abundant proteins tend to have more unique peptides identified. To understand the relative contribution of each factor, we performed rank regression by using the individual factors as the explanatory variables and the ranks of the proteomic replicates as the response variable (Methods). We found in all cases that a model including all four factors performed better than a model including only the best individual factor, suggesting that reproducibility can best be explained by a combination of factors (Figure S5D).

The factors above all contribute to protein-protein reproducibility, raising the question of whether they themselves might be sufficient to explain variation in mRNA-protein correlation. To assess this, we performed linear regression with these factors (abundance, variance, unique peptides and protein half-lives) as explanatory variables and the mRNA-protein correlation of each of the 13 different studies as response variables. We found that a combined model of the factors explained ~3-17% of the variation in mRNA-protein correlation of the different studies (Figure S6). However, the aggregated protein reproducibility explains a considerably higher percentage of the variation in mRNA-protein correlation in 12 of 13 studies. The GTEx study is the lone exception, likely a result of the independent variables (protein abundance, variance, number of unique peptides) being calculated from the GTEx study itself (Figure S6).

### Transcriptomic reproducibility also contributes to the variance in mRNA-protein correlation

Thus far, we have primarily focussed on understanding the influence of protein quantification reproducibility on mRNA-protein correlation. However, it is also likely that the reproducibility of mRNA measurements is an important factor in determining mRNA-protein correlations. To assess the impact of transcriptomic reproducibility on mRNA-protein correlation we compared transcriptomic profiles for 382 cancer cell lines from the CCLE (Ghandi et al., 2019) to those generated in a separate profiling effort (Klijn et al., 2015). We find that the median gene-wise Spearman correlation across studies was 0.75 (Methods; Figure 6A). Again this varied significantly across genes, ranging from −0.05 to 0.96.

**Figure 6.**
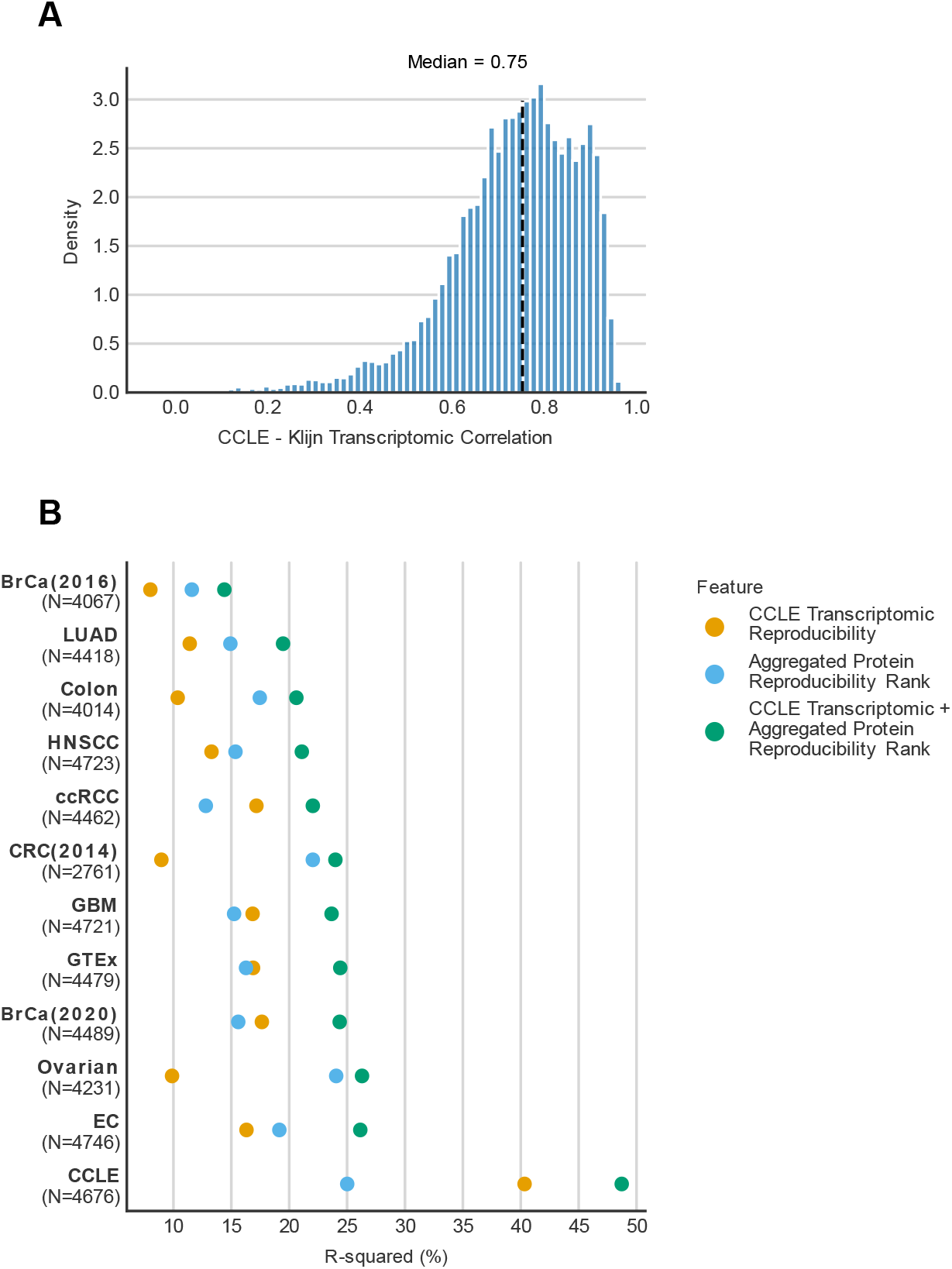
Transcriptomic reproducibility contributes to the variance in mRNA-protein correlation. (A) Histogram showing the distribution of the gene-wise correlation between experimental transcriptomic replicates of 382 cancer cell lines. The black line represents the median. (B) For each of the 13 studies analysed here, the R-squared obtained by regressing mRNA-protein correlation on transcriptomic reproducibility and aggregated protein reproducibility scores individually and in combination over the same set of proteins is shown in the dot plot. The number of proteins analysed for each study is indicated in brackets below the study on the Y-axis.

We used a linear regression model to quantify, in all thirteen proteogenomic studies, how much of the variation in mRNA-protein correlation could be explained by transcriptomic reproducibility. We found that the median variance explained was 15%. In most studies (8/13), our aggregated protein reproducibility explained a higher proportion of the variance than the mRNA reproducibility (Figure 6B).

Compared to the other studies, the CCLE study had a strikingly higher percentage of variance explained by transcriptomic reproducibility (40%). This is presumably because there is a large overlap in the set of samples used to compute the transcriptomic reproducibility and the CCLE mRNA-protein correlation unlike the other studies. For the CCLE the variance explained by mRNA-mRNA reproducibility is higher than the variance explained by protein-protein reproducibility. However, the mRNA-mRNA reproducibility was estimated using a much higher number of cell lines (382 vs 18 for protein-protein reproducibility) which we reasoned could explain the increased variance explained. To test this hypothesis we downsampled the available transcriptomic data to make the comparison more equal (sampling 18 cell lines with transcriptomes at random, Methods). We found that using this approach the protein-protein reproducibility explained more of the mRNA-protein variability than the mRNA-mRNA reproducibility (on average ~2.8 times). This suggests that protein-protein reproducibility may influence mRNA-protein correlation more than mRNA-mRNA reproducibility does, but that 18 cell lines is not sufficient to obtain a robust estimate of protein-protein reproducibility.

The Spearman correlation between aggregated protein reproducibility and CCLE transcriptomic reproducibility is 0.37 across 4,795 proteins. This suggests that there is some agreement between the reproducibility of proteins and transcripts, and that to some extent proteins that are reproducibly measured are encoded by transcripts that are more reproducibly measured. To assess if both mRNA and protein reproducibility *independently* contribute to the variability of mRNA-protein correlation across all 13 studies we used a linear model with the two factors as independent variables and mRNA-protein correlation as the dependent variable. We found that in all cases the two factors together explained a higher proportion of variance than either factor alone (p-value < 0.001, Likelihood ratio test). In the case of the CCLE study (used to calculate the mRNA reproducibility and one of the three studies used to calculate protein reproducibility) the two factors together explained 48% of the variance. For the 12 other studies the two factors together explained ~14-26% of the variance (Figure 6B). These observations suggest that the reproducibility in transcriptomic and proteomic data contribute strongly and somewhat independently to the variability observed in mRNA-protein correlation.

### Metabolic pathways with higher than average mRNA-protein correlations may reflect differential reproducibility rather than differential post-transcriptional regulation

Previous work has found that certain pathways and processes are enriched in proteins that have higher or lower than average mRNA-protein correlation. For instance, ribosomal subunits have been found to have consistently lower than average mRNA-protein correlations across multiple studies (Clark et al., 2019; Mertins et al., 2016; Zhang et al., 2014, 2016) while members of pathways related to amino-acid metabolism have been found to have higher than average mRNA-protein correlation (Clark et al., 2019; Huang et al., 2021; Jarnuczak et al., 2021; Mertins et al., 2016; Zhang et al., 2014, 2016). This variation across functional groups has been attributed to differential post-transcriptional regulation. However our observation that both protein-protein measurement reproducibility and mRNA-mRNA measurement reproducibility contribute significantly to the variation in mRNA-protein correlation across genes suggests an alternative explanation – some pathways may have higher or lower than average mRNA-protein correlations simply because their component proteins are more reproducibly measured. To test this hypothesis we first performed pathway enrichment analysis on the mRNA-protein correlations from the CCLE and ovarian datasets (Methods; Figure 7 and S6). Consistent with previous studies we observed that proteins with high mRNA-protein correlation are enriched in gene sets involved in environmental information processing (e.g. ECM-receptor interaction, Cell adhesion molecules) and metabolic pathways (e.g. Arginine and proline metabolism, Glutathione metabolism, Purine metabolism), while proteins with low mRNA-protein correlation are enriched in annotations related to housekeeping protein complexes (e.g. spliceosome, ribosome, RNA polymerase) (Figure 7; Table S3 and S4). To assess whether these enrichments could simply be attributed to variable reproducibility, we next performed pathway enrichment analysis on the CCLE and ovarian mRNA-protein correlation data *after* accounting for variation in protein-protein and mRNA-mRNA reproducibility (Methods). We found that the ‘housekeeping’ protein complexes were still identified as being enriched among proteins with lower than average mRNA-protein correlation but that the metabolic pathways were no longer enriched in proteins with higher than average mRNA-protein correlation in both CCLE and ovarian studies (Figure 7 and S7, Table S3 and S4). Other pathways with higher than average mRNA-protein correlation related to environmental information processing were also no longer significant after adjusting for reproducibility. This suggests that while large housekeeping protein complexes such as the ribosome have lower than average mRNA-protein correlation that may be attributed to post-transcriptional mechanisms, the higher-than-average mRNA-protein correlation previously observed for metabolic pathways may simply reflect more reproducible measurements of their constituent proteins and transcripts.

**Figure 7.**
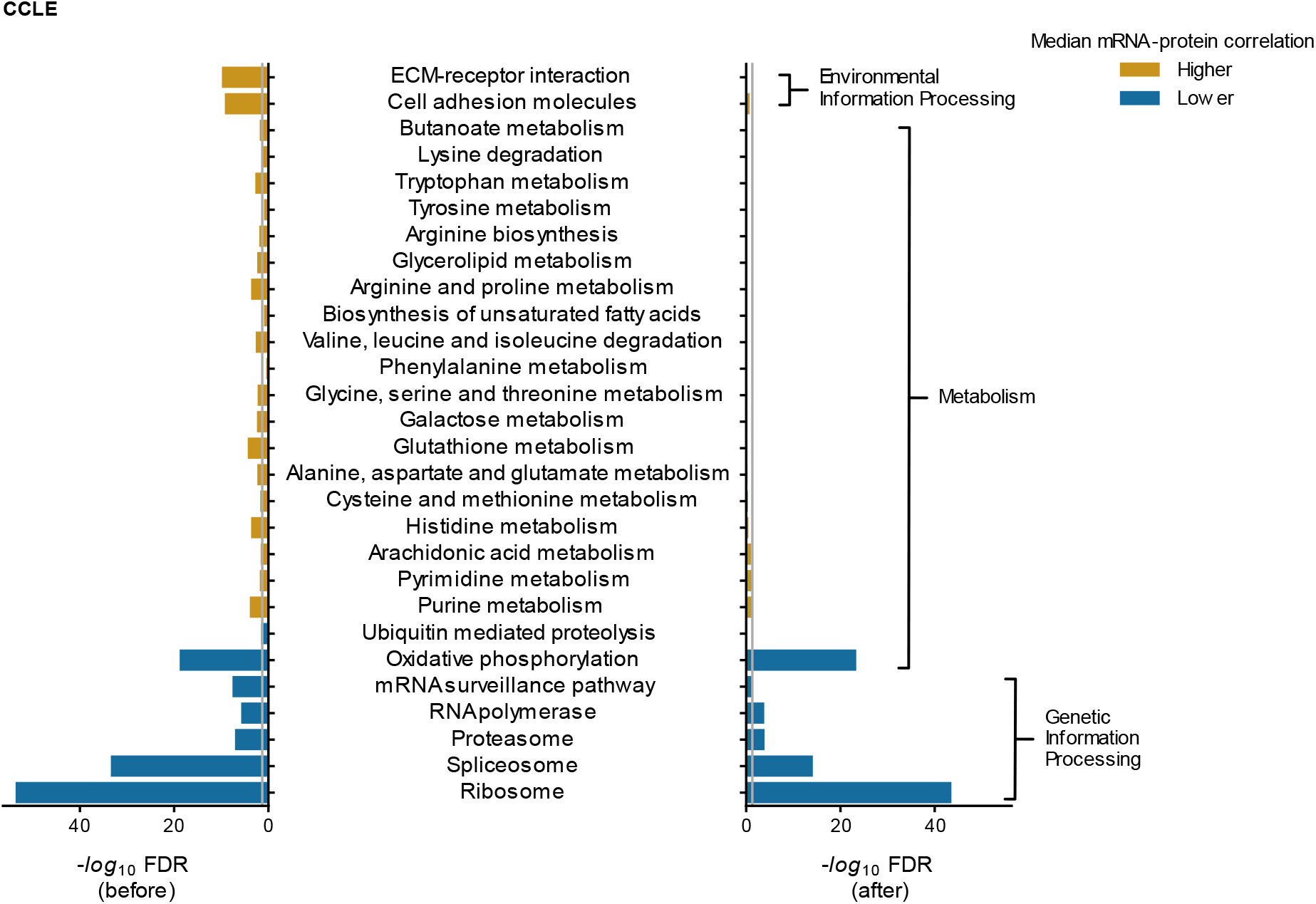
Metabolic pathways with higher than average mRNA-protein correlations may reflect differential reproducibility. Bar charts displaying the KEGG pathway enrichment analysis of the CCLE mRNA-protein correlation before (left) and after (right) accounting for protein-protein and mRNA-mRNA reproducibility. The −log_10_ of Benjamini-Hochberg FDR corrected p-values calculated using Mann-Whitney U test is deemed as enrichment for the pathway. For each bar chart, the grey line indicates the threshold considered for significant enrichment (FDR < 0.05). If the enrichment is below the threshold, then it is not considered significant. The bars are coloured orange if the median mRNA-protein correlation of genes within the pathway > median mRNA-protein correlation of genes *not* in the pathway, otherwise the bars are coloured blue.

## Discussion

Tumour proteogenomic studies have identified the correlation between mRNA and protein abundances to be only moderate (~0.4). A consistent observation from these studies is that some functional groups, such as the ribosome, have lower-than-average mRNA-protein correlation while others, such as pathways involved in amino-acid metabolism, have higher than average mRNA-protein correlation. Much of this variation in the mRNA-protein correlation has been attributed to post-transcriptional regulation. Here, we have demonstrated that the reproducibility of protein and transcript measurements is a very significant factor. We found that the observed mRNA-protein correlation is very dependent on protein reproducibility and that more reproducibly measured proteins display higher mRNA-protein correlation. After taking this into account we found that some pathways previously identified as having a high mRNA-protein correlation are likely just more reproducibly measured. We therefore suggest that conclusions about functional groups with higher or lower mRNA-protein correlation, especially with regard to the potential role played by post-transcriptional regulation, should be made only after accounting for variation in the measurement reproducibility of their constituent proteins. To this end, by integrating multiple studies, we have developed a rank aggregation method which we used to generate an aggregate protein reproducibility rank across all proteomic replicate studies for each protein. This aggregate protein reproducibility rank can explain a significant amount of the variance across multiple proteogenomic studies and may be useful for identifying those proteins that can be reliably and reproducibly measured by mass spectrometry. Such proteins may be more useful to assay in, e.g. diagnostic panels.

Recently there have been a number of attempts to predict protein abundances from transcriptomic data that have achieved modest success (Barzine et al., 2020; Fortelny et al., 2017; Li et al., 2019; Yang et al., 2020). We found here that proteins that are more reproducibly measured across experimental replicates are better predicted using machine learning. This suggests that one of the factors limiting the accuracy of machine learning methods to predict protein abundances is that the protein abundance measurements themselves are not reproducible. It may therefore be worth evaluating future methods on the subset of proteins that can be reproducibly measured.

Our emphasis here has been on understanding how variability in the measurements of individual proteins can influence the mRNA-protein correlations observed in published tumour proteogenomic studies. We have shown that proteins/transcripts that are more reproducibly measured tend to have higher mRNA-protein correlation and we have identified a number of factors (e.g. protein abundance) that influence variation in measurement reproducibility. There are of course additional factors that influence the *global* reproducibility of proteomes and transcriptomes quantified from ‘replicates’ of the same sample. These include real biological variation (e.g. tumour heterogeneity resulting in two samples of the same tumour having different profiles) and technical variation (e.g. variation in sample preparation between different runs of the same sample). We have not here been able to address how much of the variance in the measurements of individual proteins can be attributed to these global factors. It is likely that reducing these sources of global variation, e.g. through automated sample preparation, will improve the overall reproducibility of protein measurements.

We note also that our analyses do not reflect the best possible reproducibility of proteomic and transcriptomic measurements, but rather they reflect the reproducibility observed in existing large-scale proteogenomic datasets. Indeed we see that more recent proteogenomic studies have higher mRNA-protein correlation, suggesting that methodological improvements are already reducing the sources of noise in these approaches.

## Methods

### Data collection

The datasets analysed were downloaded from the links provided in the key resources table

For studies (Clark et al., 2019; Dou et al., 2020a; Gillette et al., 2020; Huang et al., 2021; Krug et al., 2020; Wang et al., 2021) both the transcriptomic and proteomic profiles were obtained from the CPTAC API (Lindgren et al., 2021). For colorectal (Zhang et al., 2014) and breast cancer (Mertins et al., 2016) studies, the transcriptomic data were downloaded from cBioPortal while proteomic data was obtained from the supplemental materials. For the ovarian cancer study (Zhang et al., 2016), the transcriptomic data were downloaded from the https://gdac.broadinstitute.org/and proteomic data from the supplemental materials. For colon cancer (Vasaikar et al., 2019), GTEX (Jiang et al., 2020) and NCI60 (Guo et al., 2019) cancer cell lines studies, both the transcriptomic and proteomic data were obtained from the supplemental tables. For CCLE study, the transcriptomic data was downloaded from the cancer dependency map portal (https://depmap.org/portal/ccle/) and proteomic data was downloaded from the supplemental materials.

### Pre-processing proteomic and transcriptomic profiles

Proteomics and transcriptomics data were obtained from the studies listed in the Resources Table. The proteomics datasets contained a considerable number of missing values, identified as NaNs in most studies or 0s in (Zhang et al., 2014). Within each study we restricted our analyses to proteins that were measured in at least 80% of samples. The same filtering was applied to transcriptomics, requiring transcripts to be measured in 80% of samples. In some datasets, multiple protein isoforms from the same gene were available, we aggregated these using the mean to calculate a ‘gene level’ summary.

The CCLE study repeatedly profiled two 10-plexes (18 cell lines) one year apart in order to assess the reproducibility of the proteomic profiling. These replicates are used to perform the assessment of the reproducibility of protein measurements presented in Figure 1. In addition to these 18 cell lines, 3 cell lines were screened in duplicate as part of standard 10-plex runs. As suggested in the CCLE guide (Nusinow and Gygi, 2020) for these three cell lines we selected the profiles which correlate best with the transcriptomic data for our analyses here.

### Computation of correlation coefficient

All data was processed through the standard pipeline described above before computing correlation. Correlation between (i) mRNA-protein, (ii) protein-protein and (iii) mRNA-mRNA was computed using the Spearman rank correlation. Considering the non-linear relationship between mRNA and protein abundances and the possibility of outliers, we choose to report median Spearman rank correlation for all studies. We note that with the exception of (Mertins et al., 2016), which used Pearson’s correlation, all published studies have also used Spearman’s correlation. For each gene in each study, samples with missing values were ignored when computing the correlation.

### Assessing proteomic and transcriptomic reproducibility

The quantitative proteomics of the CCLE (Nusinow et al., 2020) data contained three replicates of the proteomic profiles. In the first year, 18 cell lines (two 10-plexes) were quantified (R1). The same cell lines were quantified twice (R2, R3) the following year. The correlation between replicates: R1-R2, R1-R3 and R2-R3 were 0.7, 0.71 and 0.88 respectively. We chose to use the R1 and R3 proteomic profiles to compute the replicate correlation as R1-R3 has the median correlation out of the three replicate pairs.

To assess the reproducibility of transcriptomic data we considered two studies that had quantified transcripts in tumour-derived cell lines. One of the studies chosen was the CCLE transcriptomic study for which we have previously assessed the mRNA-protein correlation. The CCLE transcriptomic study (Ghandi et al., 2019) had profiled 1076 and (Klijn et al., 2015) had profiled 675 cancer cell lines using RNA-Seq. These two studies had quantified the transcripts in different labs in different years. However, the two studies had 382 cell lines and 13,226 genes in common. The transcriptomic reproducibility was computed using the Spearman rank correlation coefficient for the transcriptomic measurements across the 382 common cell lines of the studies. The standard pipeline for pre-processing was applied before assessing the reproducibility of the transcriptomic studies.

While the CCLE transcriptomic reproducibility was computed using 382 cell lines, the CCLE proteomic reproducibility was computed using 18 cell lines only. The common cell lines between the transcriptomic and proteomic replicates was <10. Therefore, to compare the predictive power of transcriptomic reproducibility and proteomic reproducibility in explaining the variation in mRNA-protein correlation of the different studies, the transcriptomic reproducibility was computed for 18 random cell lines over 100 iterations. The mean R^2^ obtained for each study was computed for estimating the predictive power for transcriptomic reproducibility.

### Computation of Deciles

Deciles were computed using the pandas qcut method. Each decile contains ~10% of the overall number of items to be stratified. In some cases, due to ties, these deciles are not uniformly sized.

### Protein complex membership

Information on protein complex membership was obtained from CORUM (Giurgiu et al., 2019) (all complexes data). A protein was marked as a protein complex subunit if it is identified in CORUM data.

### Protein half-lives

The half-lives of proteins were obtained from (Zecha et al., 2018) study. The median half-life of all proteins from the list was computed. Proteins with half-lives > median were encoded to have ‘long’ half-life while the others were encoded to have ‘short’ half-life.

### Rank Aggregation

For each of the three proteomic studies with replicates (ovarian, colon, CCLE) ranks were assigned based on increasing correlation and normalized by dividing over the total number of proteins in the dataset. Only proteins that were measured in 2 out of the 3 datasets were considered for the aggregated list. For proteins measured in only 2 studies, we imputed the third normalised rank as 0.5. For all proteins, we then computed the mean rank that has been considered to be the aggregated rank of the protein (Figure S2A).

We compared the aggregated list of proteins obtained through our method of aggregation (Figure S2A) with other aggregated lists which we calculated using other algorithms - robust rank aggregation (Kolde et al., 2012), Stuart (Stuart et al., 2003), BordaFuse (Aslam and Montague, 2001) and, Markov Chain Aggregator (MC4) (Dwork et al., 2001). To assess the performance of different aggregation methods, we used linear models wherein the mRNA-protein correlation of the three studies containing replicate proteomic profiles was regressed on the different aggregated lists of protein reproducibility. The aggregated list using our method could best explain the variation in mRNA-protein correlation in the colorectal cancer and CCLE studies, while the BordaFuse method best explained the variation in the ovarian cancer study (Figure S2B), followed by our approach. As our ‘average normalized rank’ approach overall has the highest R-squared, we chose this method to aggregate the correlations of proteomic replicate profiles.

Notably, the aggregated ranks computed incorporating the imputation method were on average 5% better in explaining mRNA-protein correlations of the different studies than the aggregated ranks computed without imputation.

### Linear Regression Models

All linear regression was carried out using the statsmodel package in Python.

#### Assessing the relationship between protein-protein reproducibility and mRNA-protein correlation (Figure S1A)

To understand the variance in mRNA-protein correlation explained by protein complex membership and protein-protein reproducibility we used three different linear models given by the equations -

- Protein complex membership only: *c* (*g*) = *α* + *β* * *pcm* (*g*)
- protein-protein reproducibility only: *c* (*g*) = *α* + *β* * *p*(*g*)
- Protein complex membership and protein-protein reproducibility:

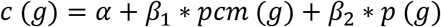

where *c* (*g*) is the mRNA-protein correlation for each gene, *pcm* (*g*) is the protein complex membership for each gene, *p*(*g*) is the protein-protein correlation for each gene and the coefficients *α*, *β*, *β*_1_ and *β*_2_ are computed using the ordinary least squares regression method. The protein complex membership is indicated as 1 if a gene is a protein complex member, else 0. For all the models, mRNA-protein correlation is assessed over the same set of genes in each study. R^2^ is used to assess the predictive power of the explanatory variables in explaining the variation of the response variable.

#### Assessing the ability of different aggregation approaches to rank protein-protein reproducibility (Figure S2B)

To identify the best aggregation method for protein-protein reproducibility, we compared the variance in mRNA-protein correlation explained by different aggregation methods using linear models given by the equations -

- Robust rank aggregation: *c* (*g*) = *α* + *β* * *P_rra_*(*g*)
- Stuart aggregation method: *c* (*g*) = *α* + *β* * *p_stuart_*(*g*)
- BordaFuse aggregation method: *c* (*g*) = *α* + *β* * *P_bf_*(*g*)
- Markov chain aggregator 4: *c* (*g*) = *α* + *β* * *P_mc4_*(*g*)
- Average normalized rank: *c* (*g*) = *α* + *β* * *p_a_*(*g*)

where *c* (*g*) is the mRNA-protein correlation for each gene, *p_rra_*(*g*), *P_stuart_*(*g*), *P_bf_*(*g*), *P_mc4_*(*g*)and *p_a_*(*g*)are the aggregated protein reproducibility ranks computed using robust rank aggregation, Stuart, BordaFuse, Markov chain aggregator 4 and average normalized ranks respectively for each gene. The coefficients *α* and *β* are computed using the ordinary least squares regression method. For all the models, mRNA-protein correlation is assessed over the same set of genes in each study. R^2^ is used to assess the predictive power of the explanatory variables in explaining the variation of the response variable.

#### Comparing the ability if aggregated rank reproducibility to predict mRNA-protein correlation compared to reproducibility calculated in individual studies (Figure S3)

For each study, we compared four different models given by the equations -

1. Ovarian protein reproducibility rank: *c* (*g*) = *α* + *β* * *P_ovarian_*(*g*)
2. CCLE protein reproducibility rank: *c* (*g*) = *α* + *β* * *P_ccie_*(*g*)
3. Colon protein reproducibility rank: *c* (*g*) = *α* + *β* * *P_coion_*(*g*)
4. Aggregated protein reproducibility rank: *c* (*g*) = *α* + *β* * *p_a_*(*g*)

where *c* (*g*)is the mRNA-protein correlation for each gene, *p_ovarian_*(*g*), *P_ccle_*(*g*), *P_colon_*(*g*)and *p_a_*(*g*) are the aggregated protein reproducibility computed using the ovarian, ccle and colon proteomic replicates individually and collectively respectively for each gene. The coefficients *α* and *β* are computed using the ordinary least squares regression method. For all the models, mRNA-protein correlation is assessed over the same set of genes in each study. R^2^ is used to assess the predictive power of the explanatory variables in explaining the variation of the response variable.

#### Assessing the impact of protein measurement reproducibility on the accuracy of machine learning prediction of protein abundance (Figure S4C)

To understand the variation in protein prediction scores that can be explained by proteinprotein reproducibility, we compared four different models on prediction scores of breast and ovarian tumour studies given by the equations -

- Ovarian protein reproducibility rank: *p_scores_*(*g*) = *α* + *β* * *p_ovarian_*(*g*)
- CCLE protein reproducibility rank: *p_scores_*(*g*) = *α* + *β* * *p_ccle_*(*g*)
- Colon protein reproducibility rank: *p_scores_*(*g*) = *α* + *β* * *p_colon_*(*g*)
- Aggregated protein reproducibility rank: *p_scores_*(*g*) = *α* + *β* * *p_a_*(*g*)

where *p_scores_*(*g*) is the prediction score that is the Pearson correlation between the predicted and actual protein abundance value obtained from the best predicting model in NCI CPTAC Proteogenomics DREAM challenge, *p_ovarian_*(*g*), *P_ccle_*(*g*), *P_colon_*(*g*) and *p_a_*(*g*) are the aggregated protein reproducibility computed using the ovarian, ccle and colon proteomic replicates individually and collectively respectively for each gene. The coefficients *α* and *β* are computed using the ordinary least squares regression method. For all the models, protein reproducibility rank is assessed over the same set of genes in each study. R^2^ is used to assess the predictive power of the explanatory variables in explaining the variation of the response variable.

#### Assessing the impact of protein abundance, protein variance, unique peptides, protein half-lives and aggregated protein reproducibility on mRNA-protein correlation (Figure S6)

To understand the variance in the mRNA-protein correlation explained by the factors (protein abundance, protein variance, unique peptides and protein half-lives) influencing protein reproducibility, we used two different linear models given by the equations -

- Other factors ():

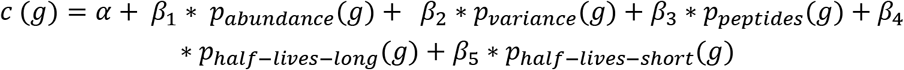
- Aggregated protein reproducibility: *c* (*g*) = *α* + *β* * *p_a_*(*g*)

where *c* (*g*) is the mRNA-protein correlation for each gene, *P_abundance_*(*g*) is the protein abundance for each gene obtained from the GTEx project, *p_variance_*(*g*) is the variance of the protein abundance for each gene obtained from the GTEx project, *p_peptides_*(*g*) is the number of unique peptides for each gene obtained from the GTEx project, *P_half-lives-long_*(*g*)and *P_half-lives-short_*(*g*) are the half-lives of each gene (long and short), *p_a_*(*g*) are the aggregated protein reproducibility computed using the ovarian, CCLE and colon proteomic replicates individually and collectively respectively for each gene and the coefficients:*α*, *β*, *β*_1_, *β*_2_, *β*_3_, *β*_4_ and *β*_5_ are computed using the ordinary least squares regression method. For all the models, mRNA-protein correlation is assessed over the same set of genes in each study. R^2^ is used to assess the predictive power of the explanatory variables in explaining the variation of the response variable.

### Rank Regression

We used rank regression to assess the contribution of various factors (protein abundance, unique peptides and protein half-lives) to explaining the variance in protein measurement reproducibility. We assessed both the aggregated ranks and the reproducibility measured in each individual study. We converted the protein reproducibility measurements from the three studies with replicates (ovarian, colon, CCLE) to ranks.

The potential factors such as protein abundance and unique peptides had a large range, therefore both the factors were log transformed and linear regression was performed.

#### Assessing the impact of protein abundance, protein variance, unique peptides, protein half-lives on the reproducibility of proteins (Figure S5D)

To understand the variance in the protein reproducibility ranks explained by the potential factors (protein abundance, protein variance, unique peptides and protein half-lives), we used four different linear models given by the equations -

- Protein abundance only: *rank* (*g*) = *α* + *β* * *P_abundance_*(*g*)
- Protein variance only: *rank* (*g*) = *α* + *β* * *P_variance_*(*g*)
- Unique peptides only: *rank* (*g*) = *α* + *β* * *p_peptides_*(*g*)
- Protein half-lives encoded as long and short:

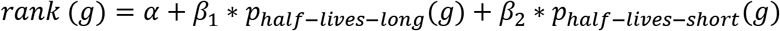
- Protein abundance, unique peptides and protein half-lives combined:*rank* (*g*) = *α* + *β*_1_ * *P_abundance_*(*g*) + *β*_2_ * *P_variance_*(*g*) + *β*_3_ * *P_peptides_*(*g*) + *β*_4_ * *P_half-lives-long_*(*g*) + *β*_5_ * *P_half–lives–short_* (*g*)

where *rank* (*g*) is the protein reproducibility rank for each gene, *P_abundance_*(*g*) is the protein abundance for each gene obtained from the GTEx project, *p_variance_*(*g*) is the variance of the protein abundance for each gene obtained from the GTEx project, *p_peptides_*(*g*) is the number of unique peptides for each gene obtained from the GTEx project, *p_haif–lives–long_*(*g*)and P_half–lives–short_(*g*) are the half-lives of each gene (long and short) and the coefficients *α*, *β*, *β*_1_, *β*_2_, *β*_3_, *β*_4_ and *β*_5_ are computed using the ordinary least squares regression method. For all the models, protein reproducibility is assessed over the same set of genes in each study. R^2^ is used to assess the predictive power of the explanatory variables in explaining the variation of the response variable.

### Pathway Enrichment Analysis

Pathway enrichment analysis was performed using the Mann-Whitney U test. Firstly, the KEGG pathways (Kanehisa et al., 2021) and their associated genes for *Homo sapiens* were downloaded using the KEGG API (https://www.kegg.jp/kegg/rest/keggapi.html). Only KEGG pathways with more than 3 genes with measured correlations were included for the enrichment analysis. The computed mRNA-protein correlations of CCLE and ovarian cancer studies were used to rank the genes. A Mann-Whitney U test was performed to assess the rank of each pathway in each dataset and p-values obtained were corrected for false discovery rate (FDR) using the Benjamini-Hochberg method. For the figures presented in Figure 7 and Figure S6 we specifically included pathways which have been previously identified as enriched in different cancer studies (Clark et al., 2019; Huang et al., 2021; Mertins et al., 2016; Zhang et al., 2014, 2016). To identify enriched pathways *after* accounting for experimental reproducibility, we regressed the CCLE and ovarian mRNA-protein correlation on both aggregated protein reproducibility ranks and mRNA-mRNA reproducibility correlations, which are based on the equations *c* (*g*) = *α* + β_1_ * *m* (*g*) + *β*_2_ * *p* (*g*), where *c* (*g*) is the mRNA-protein correlation, *m* (*g*) is the mRNA-mRNA reproducibility and *p* (*g*) is the protein-protein reproducibility and the coefficients *α*, *β*_1_ and *β*_2_ are computed based on the ordinary least squares regression method. The residuals obtained from the regression were used to rank the genes in pathway enrichment analysis. The top level categories (e.g. Metabolism, Genetic Information Processing) of the pathways were obtained from KEGG and are used to annotate the pathways in Figure 7 and S6.

### Quantification and statistical analysis

Statistical analysis is described in the method details and was carried out using Python 3.8, Pandas 1.2.5 (McKinney and Others, 2011), numpy 1.20.2 (Harris et al., 2020), SciPy 1.7.1 (Virtanen et al., 2020) and StatsModels 0.12.2 (Seabold and Perktold, 2010). The figures were created with Matplotlib 3.3.4 (Hunter, 2007) and Seaborn 0.11.1 (Waskom et al., 2020).

## Code availability

- All the notebooks containing the code, including all statistical analysis, for this project are available at -https://github.com/cancergenetics/limitations_of_omics_reproducibility

## Acknowledgments

S.R.U was funded through the School of Computer Science, University College Dublin and C.J.R was funded by the Irish Research Council Laureate Awards 2017/2018. We thank Dr. Dirk Fey, Dr. Giorgio Oliviero, Dr. Luis Iglesias Martinez and members of the Ryan lab for careful reading of the manuscript and helpful feedback. We also thank Dr. Theodoros Roumeliotis for suggesting protein variance as a factor influencing protein reproducibility.

## Supplemental Files

**Table S1.** Computed correlation between mRNA and protein measurements using the standardised pipeline. Additional details of Transcriptomic and Proteomic Data.

**Table S2.** Computed correlation between experimental proteomic replicates of ovarian and colon tumour samples and cancer cell lines using the standard pipeline and the aggregated normalized ranks.

**Table S3.** KEGG pathway enrichment analysis for CCLE study

**Table S4.** KEGG pathway enrichment analysis for ovarian cancer study

## Supplemental Figures

**Figure S1.**
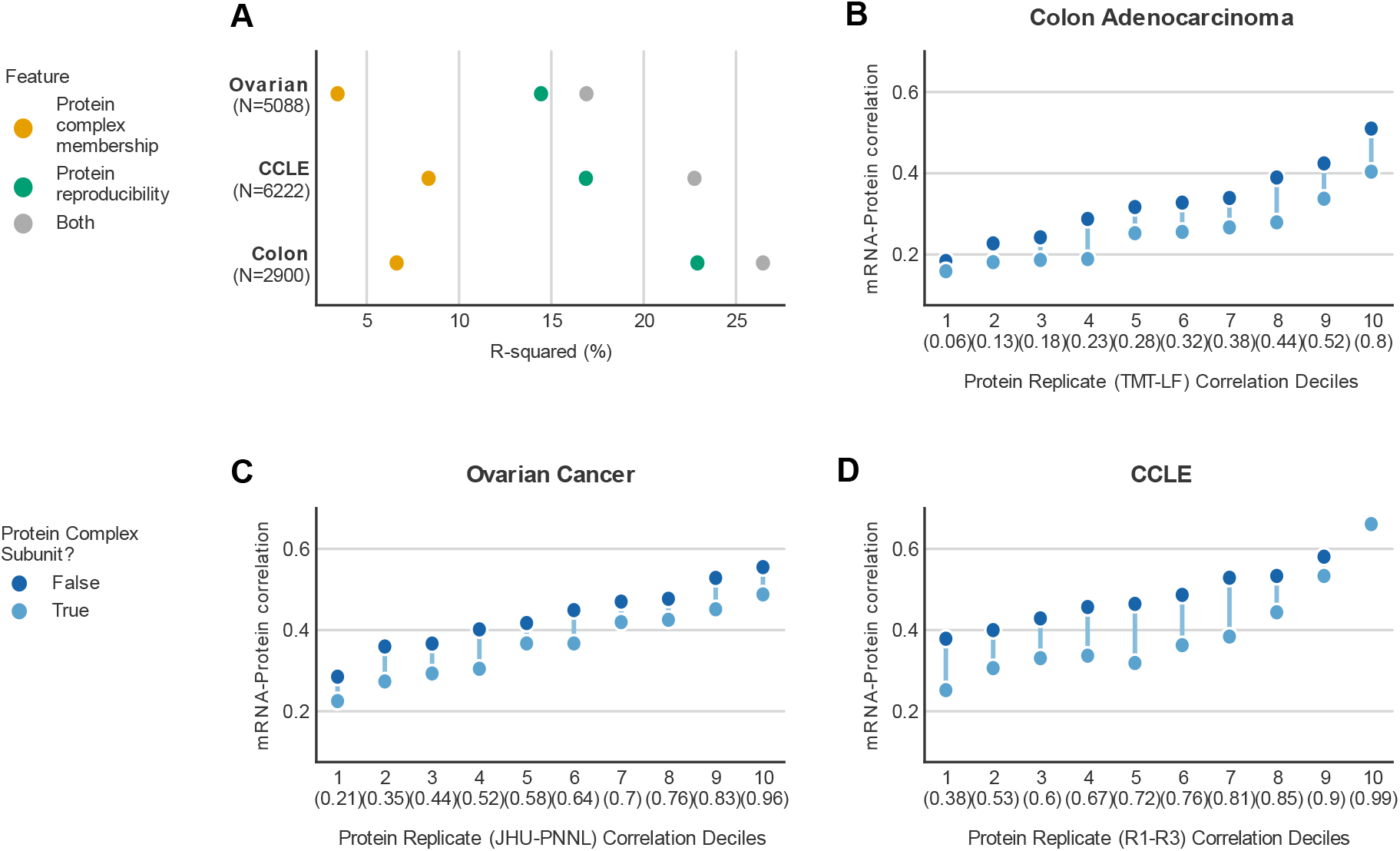
Protein complex membership and protein reproducibility contribute to the variation in mRNA-protein correlation, Related to Figure 2. (A) Dot plot displaying the R-squared obtained from regressing mRNA-protein correlation of the indicated studies on protein complex membership and their corresponding protein reproducibility over the same set of proteins. The number of proteins considered for each analysis is specified in parentheses below the study on Y-axis. (B-D) Ranged dot plots showing the mean of mRNA-protein correlation for proteins that are complex subunits (light blue dot) or not (dark blue dot) within every decile of the proteomic replicates’ correlation for colon (B) and ovarian tumour (C) and CCLE studies (D). The line represents the difference in the mean of the mRNA-protein correlation between the groups of proteins belonging to the same decile. X-axis indicates the decile number and contains the maximum correlation between the experimental proteomic replicates for that decile in parentheses.

**Figure S2.**
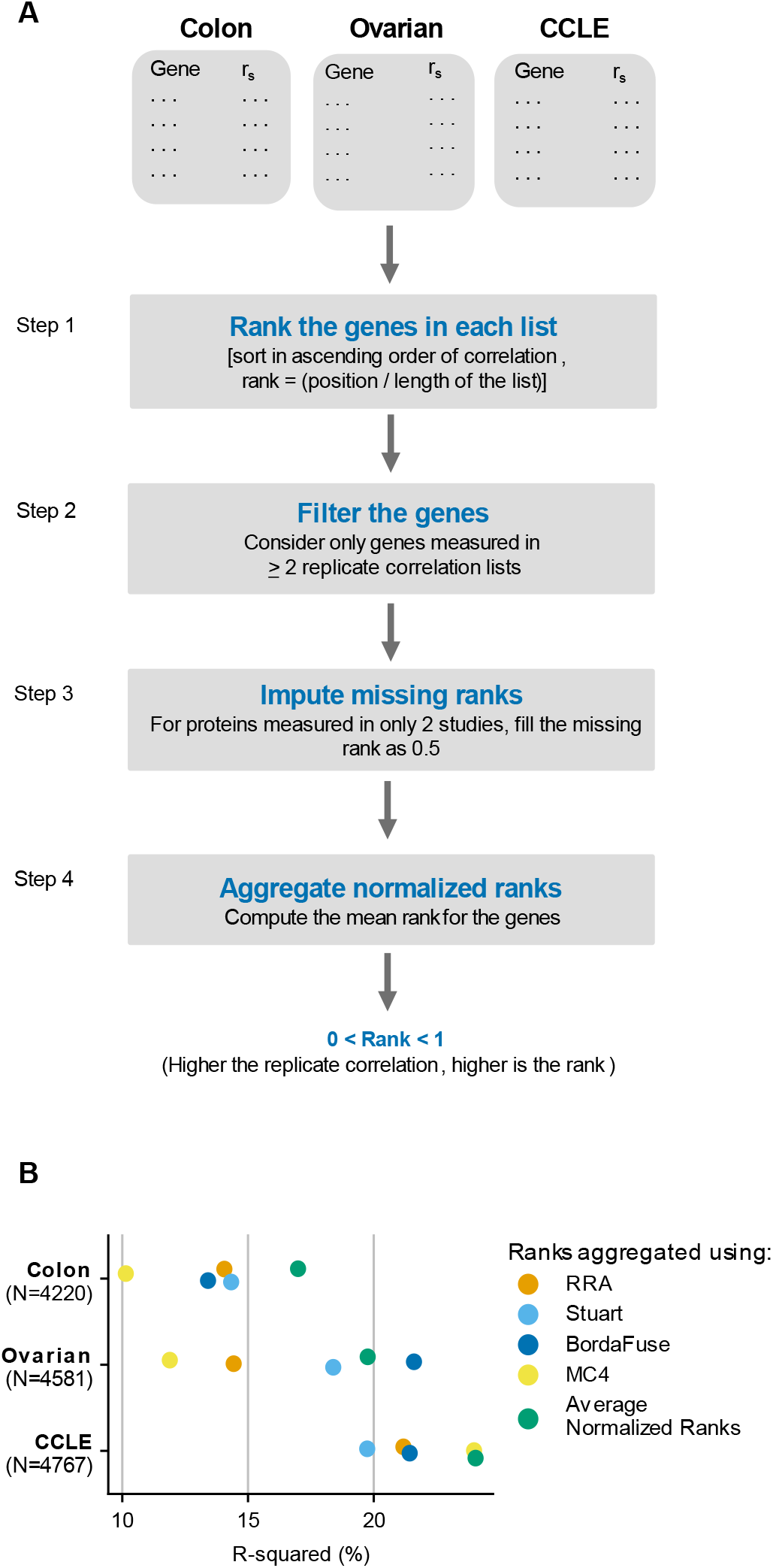
Aggregate protein reproducibility, Related to Figure 4. (A) Workflow of our computational approach to aggregate the ranks of the correlation of experimental proteomic replicates from 3 different datasets - colon, ovarian and CCLE. The computed ranks lie between zero and one. The higher the correlation between the experimental proteomic replicates, the higher the rank. (B) Dot plot displaying R-squared obtained from regressing mRNA-protein correlation of the indicated studies on the aggregated protein ranks obtained from different algorithms (robust rank aggregation, Stuart, BordaFuse, Markov chain aggregator and our method of average normalized rank) over the same set of proteins. The number of proteins considered for each analysis is specified in parentheses below the study on Y-axis.

**Figure S3.**
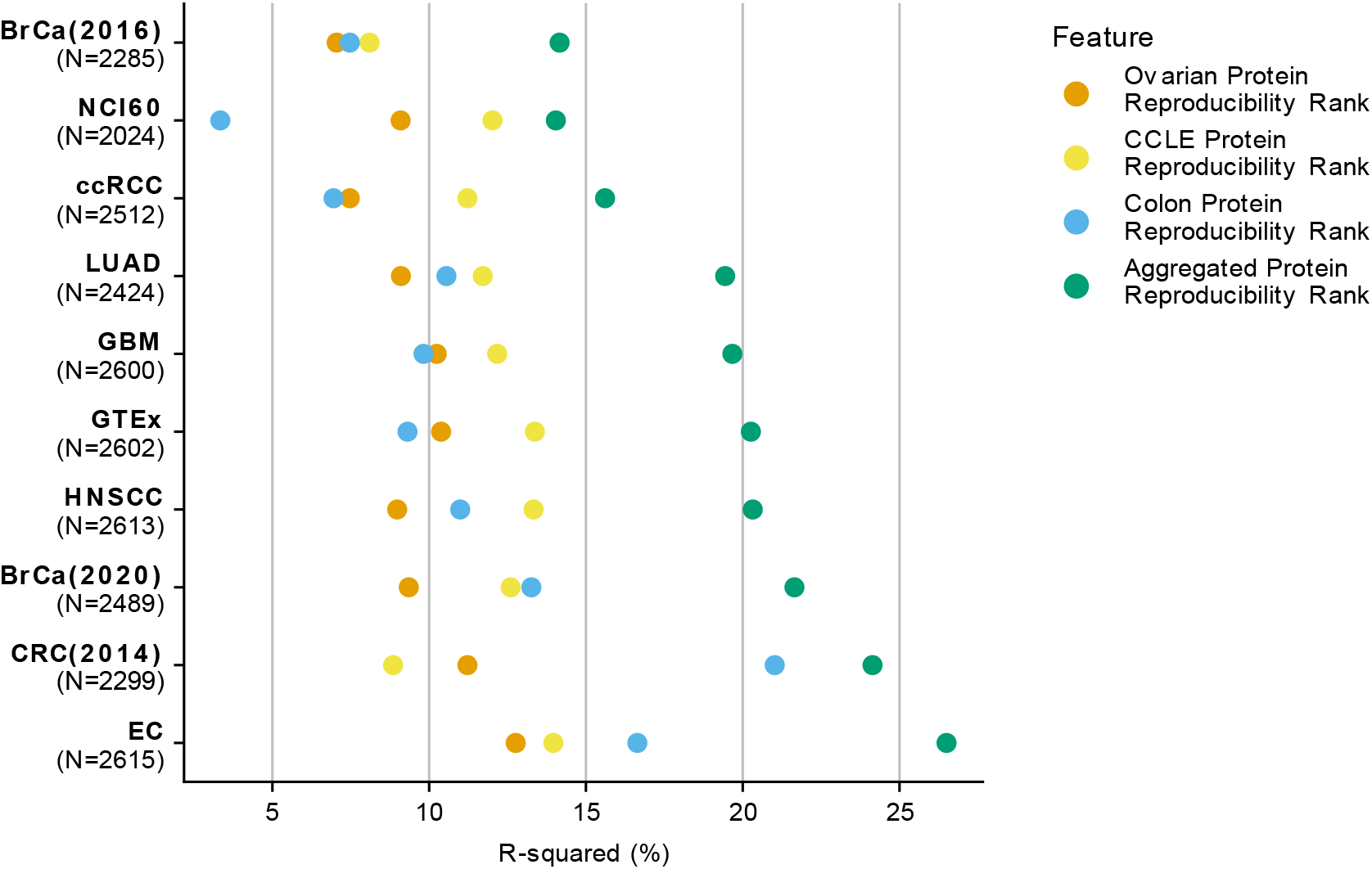
Aggregate protein reproducibility outperforms the individual protein reproducibility ranks in explaining the variation in mRNA-protein correlation, Related to Figure 4. Dot plot comparing the R-squared values obtained from regressing mRNA-protein correlation on the individual protein reproducibility ranks and the aggregated protein reproducibility rank over the same set of proteins. The number of proteins considered for each analysis is specified in parentheses below the study on Y-axis.

**Figure S4.**
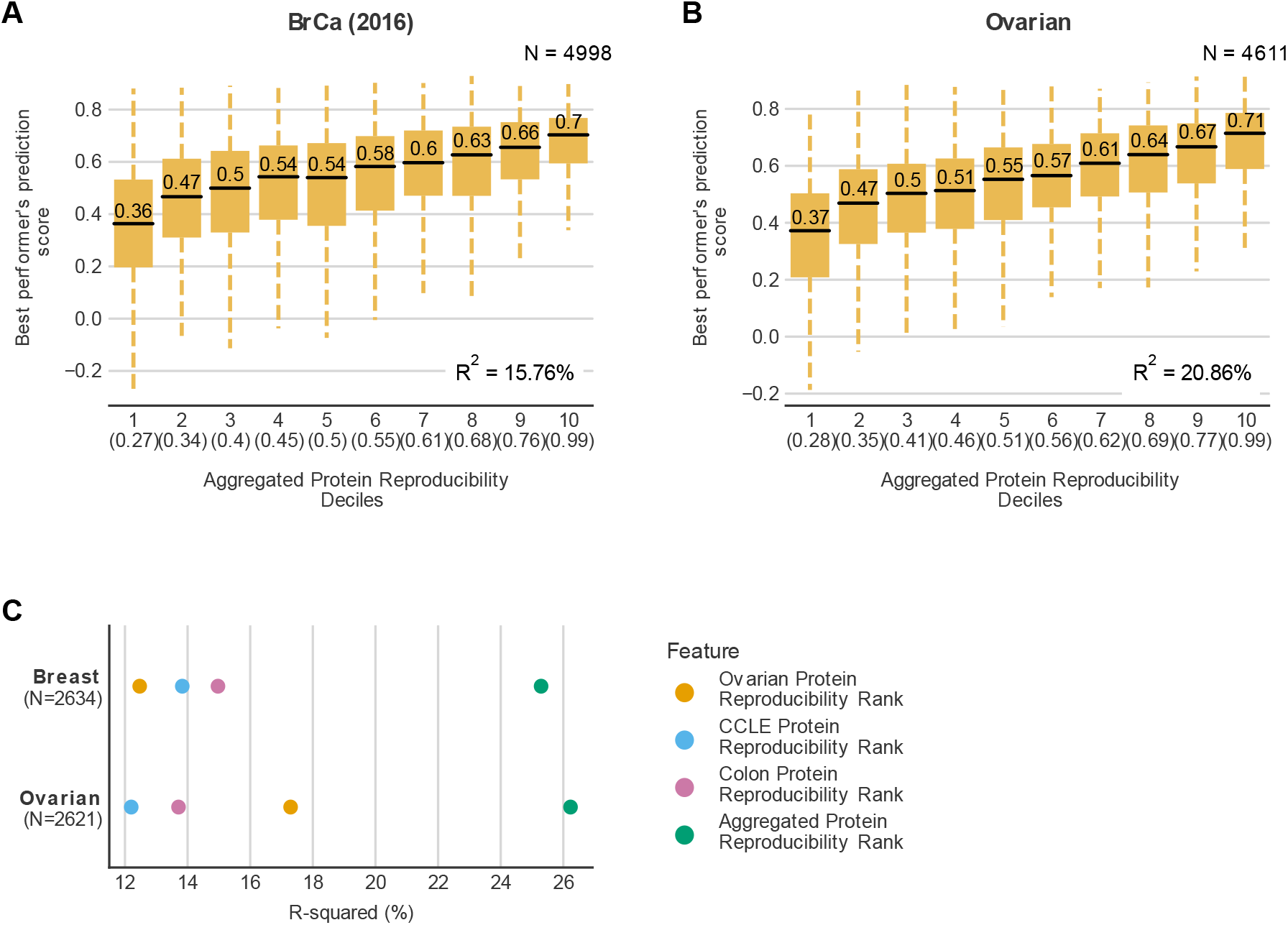
Proteins that are highly reproducible can be better predicted using machine learning, Related to Figure 4. Boxplots showing the distribution of prediction scores from the best performing model in the NCI CPTAC DREAM Proteogenomics challenge for proteins in each decile of the aggregated protein reproducibility ranks in breast (A) and ovarian studies (B). The prediction score is the Pearson correlation between the observed and predicted protein abundance. The decile is indicated on the X-axis along with the highest score of the aggregated protein reproducibility rank present within that decile. For each box plot, the black central line represents the median, the top and bottom lines represent the 1st and 3rd quartile, and the whiskers extend to 1.5 times the interquartile range past the box. Outliers are not shown. The median of each decile is indicated above the black central line for each box plot. The R^2^ obtained from regressing the prediction score on the aggregated protein reproducibility ranks is in the bottom-right corner. (C) Dot plot comparing the R-squared values obtained from regressing protein abundance prediction scores of breast and ovarian tumour studies, obtained from NCI CPTAC Proteogenomics DREAM Challenge, on the individual protein reproducibility ranks and the aggregated protein reproducibility rank over the same set of proteins. The number of proteins considered for each analysis is specified in parentheses below the study on Y-axis.

**Figure S5.**
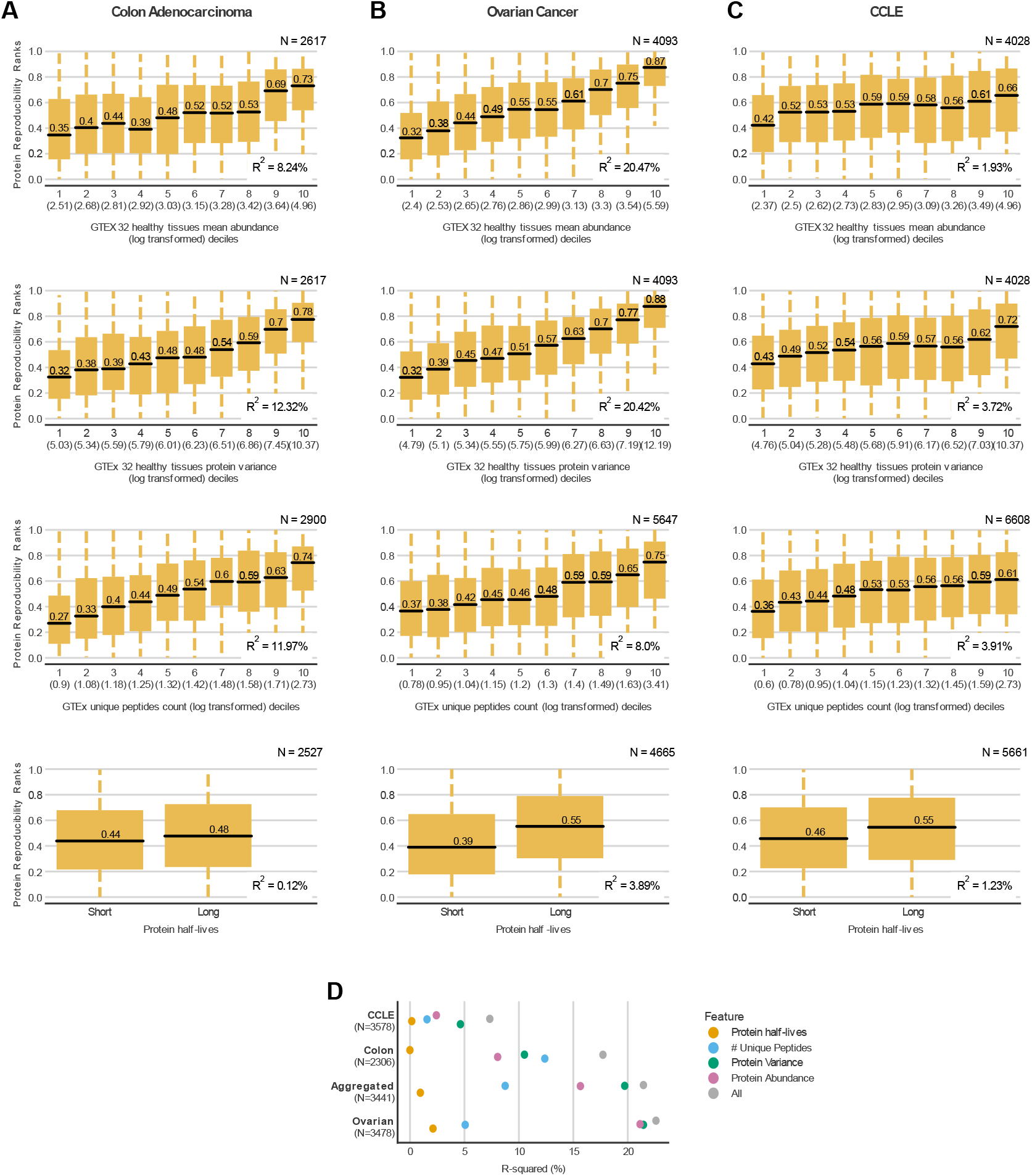
Potential factors influencing protein reproducibility in individual studies with experimental proteomic replicates, Related to Figure 5. (A-C) Similar to Fig. 6 but for the individual protein reproducibility ranks from each study. (D) Dot plot comparing the R-squared values obtained from regressing the individual protein reproducibility ranks and the aggregated protein reproducibility ranks on the potential factors that affect protein reproducibility - protein abundance, unique peptides, protein half-lives individually and all of them collectively over the same set of proteins. The number of proteins considered for each analysis is specified in parentheses below the study on Y-axis.

**Figure S6.**
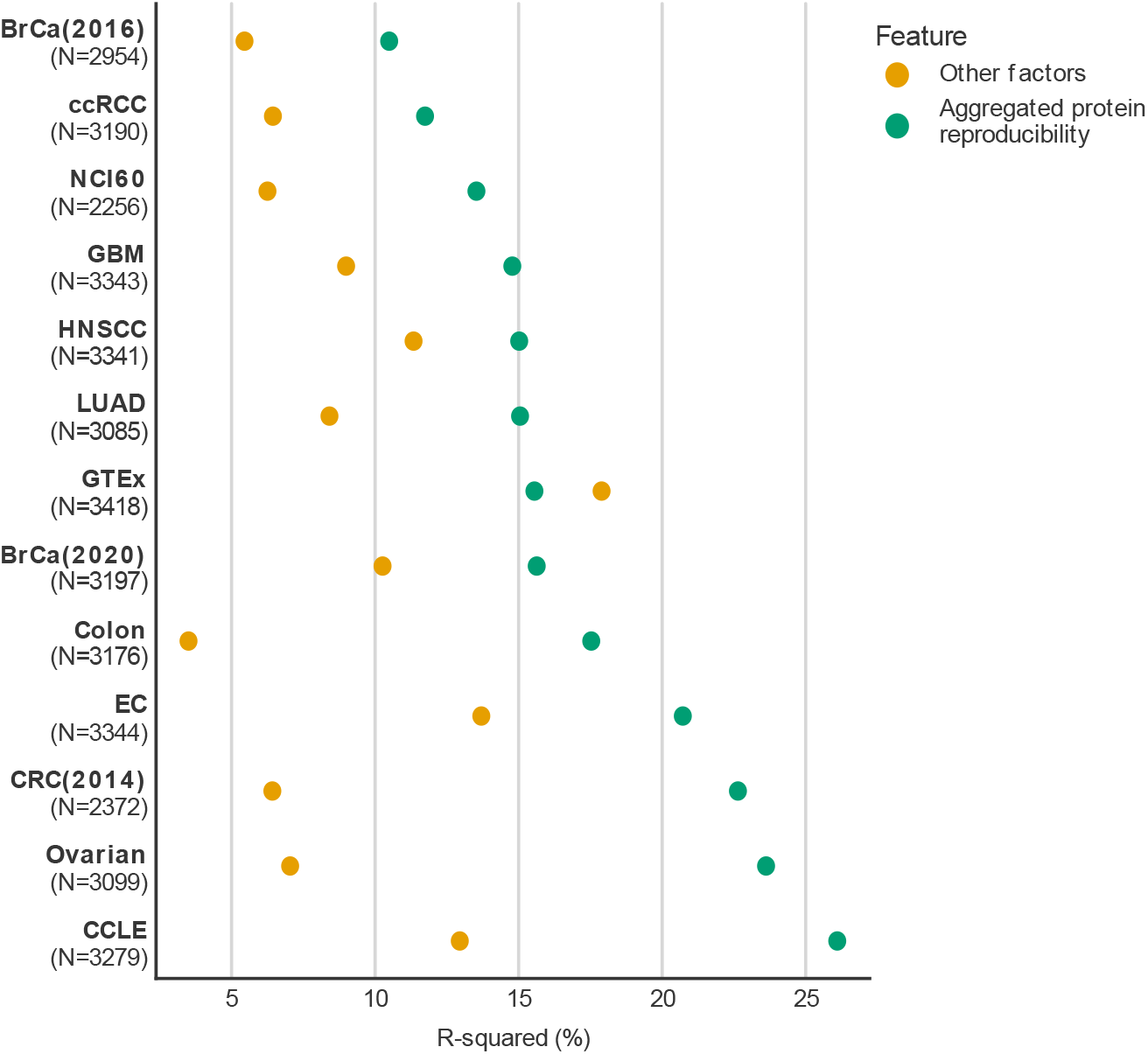
Protein reproducibility explains the variation in mRNA-protein correlation better than the other factors (protein abundance, protein variance, unique peptides, protein half-life), Related to Figure 5. Dot plot comparing the R-squared values obtained from regressing mRNA-protein correlation of the studies on the aggregated protein reproducibility ranks and the potential factors that affect protein reproducibility - protein abundance, unique peptides, protein half-lives individually over the same set of proteins. The number of proteins considered for each analysis is specified in parentheses below the study on Y-axis.

**Figure S7.**
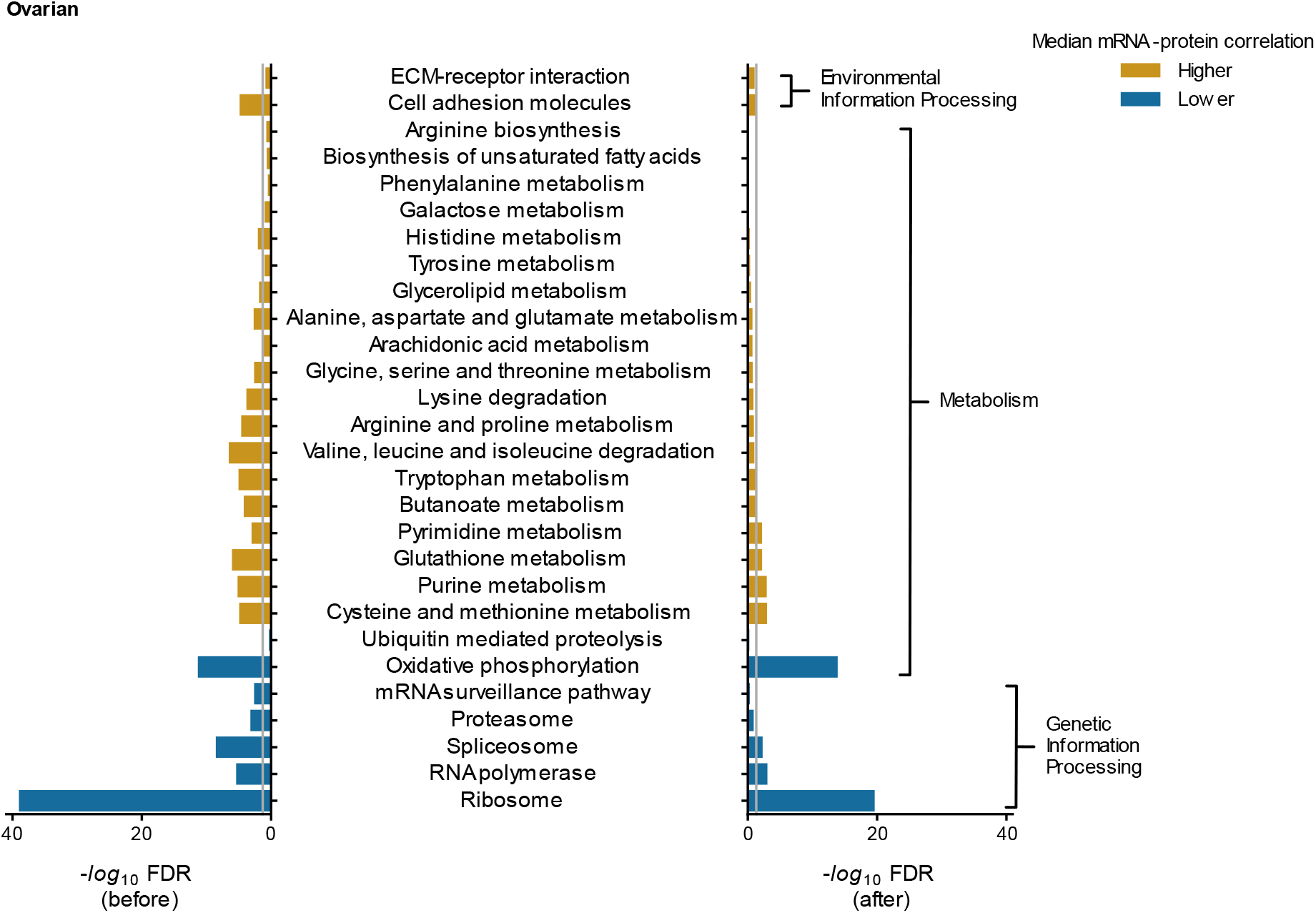
KEGG pathways enrichment analysis for ovarian cancer study, Related to Figure 7. Bar charts displaying the KEGG pathway enrichment analysis of the ovarian cancer study mRNA-protein correlation before (left) and after (right) accounting for protein-protein and mRNA-mRNA reproducibility. The −log_10_ of Benjamini-Hochberg FDR corrected p-values calculated using Mann-Whitney U test is deemed as enrichment for the pathway. For each bar chart, the grey line indicates the threshold considered for significant enrichment (FDR < 0.05). If the enrichment is below the threshold, then it is not considered significant. The bars are coloured orange if the median mRNA-protein correlation of genes within the pathway > median mRNA-protein correlation of genes *not* in the pathway, otherwise the bars are coloured blue.

